# Cryo-EM structure of the calcium release-activated calcium channel Orai in an open conformation

**DOI:** 10.1101/2020.09.03.281964

**Authors:** Xiaowei Hou, Ian R. Outhwaite, Leanne Pedi, Stephen Barstow Long

**Author notes:** Equal contribution (Xiaowei Hou, Ian R. Outhwaite). Correspondence and requests for materials should be addressed to S.B.L.

## Abstract

The calcium release-activated calcium channel Orai regulates Ca^2+^ entry into non-excitable cells and is required for proper immune function. The channel typically opens following the release of Ca^2+^ from the endoplasmic reticulum. Certain pathologic mutations render the channel constitutively open. Here, using one such mutation (H206A), we present a cryo-EM structure of Orai from *Drosophila melanogaster* in an open conformation at 3.3 Å resolution. Comparison with previous closed structures reveals that opening occurs through the outward movements of M1 helices that dilate the central pore. Repositionings of a ring of phenylalanine residues (F171) expose previously shielded glycine residues (G170) to the channel pore, despite the absence of significant rotational movement of the associated pore-lining helices. This phenylalanine ring and two rings of flanking hydrophobic amino acids act as a hydrophobic gate to control ion permeation. Extracellular M1-M2 turrets, not evident from previous Orai structures, form an electronegative pore entrance.

**Single sentence summary:** A structure of the Ca^2+^ channel Orai in an open conformation provides insight into the opening mechanism of the channel and its role in regulating selective Ca^2+^ entry into immune and other non-excitable cells.

## Introduction

Intracellular calcium signals in most non-excitable metazoan cells are augmented and shaped by the influx of extracellular Ca^2+^ ions due to opening of the calcium release-activated calcium (CRAC) channel Orai in the plasma membrane (Hogan, Lewis, & Rao, 2010). Ca^2+^ influx though the channel is necessary for activation of immune response genes in T cells and a range of other physiological processes (Feske, Prakriya, Rao, & Lewis, 2005; Lacruz & Feske, 2015; Prakriya & Lewis, 2015). Through an unusual mechanism, the depletion of Ca^2+^ from the endoplasmic reticulum (ER), rather than an increase in the cytosolic Ca^2+^ concentration, initiates opening of the channel (reviewed in (Hogan et al., 2010)). Although Ca^2+^ entry through the channel is a major contributor to intracellular Ca^2+^ signaling, the physiological functions of the channel are less appreciated than Ca^2+^ release from the ER, partly because the molecular components of the CRAC channel, the Orai protein and its regulator STIM, were identified fairly recently. Orai is an integral membrane protein that forms the pore of the channel in the plasma membrane (Feske et al., 2006; Vig et al., 2006; Yeromin et al., 2006; Shenyuan L Zhang et al., 2006). There are three Orai proteins in humans (Orai1-3). *Drosophila melanogaster* contains one ortholog (Orai), which shares 73% sequence identity to human Orai1 in the transmembrane region and is the most studied non-human Orai channel. STIM proteins (STIM1 and STIM2 in humans) are single-pass membrane proteins located in the membrane of the ER, and following Ca^2+^ depletion from the ER regulate Orai channel function (Roos et al., 2005; S. L. Zhang et al., 2005). Although the molecular mechanisms of this process are not yet fully resolved, a scheme for channel activation is becoming clear (reviewed in (Krizova, Maltan, & Derler, 2019; Lunz, Romanin, & Frischauf, 2019; Qiu & Lewis, 2019)). The “inside-out” signaling from the ER to the plasma membrane occurs at cellular locations where the ER and plasma membranes are close together (separated by ∼ 10-20 nm). Ca^2+^ release from the ER into the cytosol, which can occur via the IP3R receptor, is detected by a luminal domain of STIM from the reduction of [Ca^2+^] in the ER. An ensuing conformational change in STIM enables its cytosolic domain to interact with Orai across the divide separating the two membranes and instigates opening of the pore of Orai. Mechanisms of the Orai-STIM interaction and subsequent channel opening remain largely unresolved at the structural level. A high-resolution three-dimensional (3D) structure of the open conformation of the pore would shed light on the mechanisms of gating, ion permeation, and Ca^2+^ selectivity in the CRAC channel.

Activated CRAC channels have exceedingly low ion conductance in comparison to most other ion channels and they are highly selective for Ca^2+^ (Lepple-Wienhues & Cahalan, 1996; Prakriya & Lewis, 2006). The unitary conductance of activated CRAC channels is so low (7-25 fS in 2-110 mM Ca^2+^) that recordings of currents from single-channels have not been feasible (Prakriya & Lewis, 2003, 2006). Both of these properties, slow ion permeation and high Ca^2+^ selectivity, are fundamental to the channel’s physiological functions to generate sustained elevations of cytosolic calcium concentrations, which among other processes activate immune response genes in T cells (Hogan et al., 2010). Mutations in Orai or STIM cause a spectrum of disorders, which are typically immunological in nature when these mutations lead to loss of channel function (Lacruz & Feske, 2015). For instance, mutation of a pore-lining residue (R91W) in Orai1 causes a severe combined immune deficiency-like disorder due to lack of functional CRAC channels in the T cells of these patients (Feske et al., 2006). In addition to loss-of-function mutations, some gain-of-function mutations have been identified that allow Orai to conduct cations without activation by STIM (reviewed in (Krizova et al., 2019)). Activating mutations of Orai1 have been associated with tubular aggregate myopathy and Stormorken syndromes (Lacruz & Feske, 2015). While many of these gain-of-function mutants have reduced ion selectivity for Ca^2+^ as indicated by an altered current-voltage relationship when channels are studied using whole cell patch clamp electrophysiology, the H134A mutation of Orai1 creates a channel that is highly selective for Ca^2+^ and exhibits a similar current-voltage relationship to that of STIM-activated Orai, which suggests that the pore adopts a similar conformation to the STIM-activated channel (Irene Frischauf et al., 2017; Krizova et al., 2019; Yeung et al., 2018). Additionally, biochemical studies, which were supported by molecular dynamics simulations, suggest that the conformation of the H134A mutant is highly similar to the naturally opened channel (Irene Frischauf et al., 2017; Yeung et al., 2018). Unlike many of the activating mutations, which are located at amino acids that directly form the walls of the ion pore, the H134A mutation of Orai1 is located on the M2 helix and not exposed to the pore (X. Hou, Burstein, & Long, 2018; Xiaowei Hou, Pedi, Diver, & Long, 2012). We have shown that purified *Drosophila* Orai with the corresponding H206A mutation (Orai_H206A_) forms a constitutively active channel when reconstituted into liposomes (X. Hou et al., 2018). Orai_H206A_ exhibits properties of the STIM-activated channel, including the ability to conduct Na^+^ in the absence of divalent cations and the abilities of Mg^2+^ and Ca^2+^ to block Na^+^ current (X. Hou et al., 2018). In the absence of structural information for STIM-activated Orai, which has proven difficult to obtain, the H206A mutation provides an experimental approach with which study the 3D structure of an opened Orai pore.

The X-ray structure of a closed conformation of *Drosophila melanogaster* Orai, referred to as the “quiescent” conformation (Xiaowei Hou et al., 2012), provided the first view of the molecular architecture of the channel. This structure, determined at 3.35 Å resolution, divulged that the channel is formed from a hexamer of Orai subunits rather than a tetrameric architecture, which was anticipated at the time. Subsequent studies have shown that human Orai1 channels also assemble and function as hexamers (Xiangyu Cai et al., 2016; Yen, Lokteva, & Lewis, 2016). The structure revealed that the channel has a single ion conduction pore along a central six-fold axis of symmetry, which would be perpendicular to the membrane in a cellular setting (Xiaowei Hou et al., 2012). Each subunit contains four transmembrane helices (M1-M4). Six M1 helices, one from each subunit, form the walls of the ion pore, which is approximately 55 Å long and narrow in the closed conformation. The M2 and M3 helices surround the M1 helices but do not contribute to the pore. The M4 helices are located at the periphery of membrane-spanning portion of the channel and helices extending from them (M4-ext helices) protrude into the cytosol (X. Hou et al., 2018; Xiaowei Hou et al., 2012). The linkages between the M1 and M2 helices were not visible in the electron density and because of this, there is some ambiguity regarding which M1 helix in the hexamer is associated with which M2 helix. The location of the linkages relative to pore suggests that they would contribute to its extracellular entrance, as has been suggested by the effects of mutations of acidic amino acids within them on the binding of pore-blocking lanthanides (Gd^3+^ and La^3+^) (Yeromin et al., 2006) and the observation that such mutations reduce Ca^2+^ influx into cells (I. Frischauf et al., 2015). Structural information regarding the M1-M2 linkage would advance our understanding of the extracellular entrance.

We previously determined a ∼ 7 Å resolution X-ray structure of Orai with the activating H206A mutation (Orai_H206A_) (X. Hou et al., 2018). The structure yielded low-resolution information regarding the positioning of α-helices in an open conformation of the channel and indicated that opening involves the dilation of the pore. However, the conformations of amino acids along the pore and throughout the channel were not resolved due to the low resolution of the diffraction data (X. Hou et al., 2018). Consequently, the structure provided limited insights into mechanisms for Ca^2+^ permeation and selectivity.

One of the main questions in the field pertains to the mechanism by which ion flow is permitted through the channel in the open state. Structural, functional and computational studies have suggested that hydrophobic amino acids within the pore (particularly Phe 99 and Val 102 in human Orai1, corresponding to Phe 171 and Val 174 in *D*. Orai, respectively) function as a dynamic “gate” that prevents ion conduction when the channel is closed and permits ion permeation when the channel is open (Irene Frischauf et al., 2017; Megumi Yamashita et al., 2017). The nature of the molecular rearrangements that open the channel and the conformations of these hydrophobic residues in the open state have been queried through functional studies of mutants and through molecular dynamics simulations (Derler et al., 2013; Irene Frischauf et al., 2017; M. Yamashita et al., 2020; Megumi Yamashita et al., 2017; Yeung et al., 2018; Yeung, Yamashita, & Prakriya, 2020), but they have not yet been addressed by high-resolution structural studies of an open conformation of the channel.

In this study, we present a cryo-electron microscopy (cryo-EM) structure of Orai_H206A_ in an open conformation at 3.3 Å resolution. The structure agrees with our previous low-resolution structure of Orai_H206A_ and provides a near-atomic view of the amino acids that line the pore in an open state. The substantially improved resolution was enabled through use of Fab antibody fragments that bind to the extracellular side of the channel. Details of the opened pore, including the positioning of a critical phenylalanine within the hydrophobic region of the pore (Phe 171, corresponding to Phe 99 in human Orai1) and conformational changes throughout Orai, provide insights into the mechanisms of ion permeation, ion selectivity and gating in the channel. The structure also reveals that the connections between the M1 and M2 helices form structured turrets that create an extracellular entrance to the pore, clarifying the relationship between transmembrane helices within Orai channel subunits and suggesting a physiological role for these ordered structures.

## Results

### Antibody development and characterization

The remarkable advances in structural biology enabled by cryo-EM require that individual protein complexes embedded in vitreous ice can be identified within micrographs and that their orientations can be accurately determined (Cheng, 2015). As a consequence, complexes greater than 200 kDa and/or those with distinctly recognizable shapes typically yield higher resolution cryo-EM structures than smaller complexes or those with fewer distinguishing features. The somewhat spherical overall shape of the Orai channel and its relatively low molecular weight (∼ 144 kDa) make obtaining a high-resolution cryo-EM reconstruction difficult. We therefore pursued a structure of Orai_H206A_ in complex with a monoclonal Fab antibody fragment in order to add shapeliness and increase mass.

The monoclonal antibody 19B5 was developed by immunizing mice with purified Orai protein and selecting antibodies that bound to the structured form of the channel but not to denatured protein (Figure 1 - figure supplement 1, Methods). The ability of the 19B5 antibody to bind to the structured channel but not to denatured protein was an important criterion because an antibody that binds to a structural epitope would be more useful for cryo-EM. Using mutagenesis and fluorescence-detection size exclusion chromatography (Kawate & Gouaux, 2006) to map its binding epitope, we determined that 19B5 binds to the extracellular side of the channel in the loop connecting the M1 and M2 helices (Methods). The properties that 19B5 binds to this loop and preferentially binds to the structured form of the channel suggested that the M1-M2 connection adopts a structured conformation even though density for this loop was weak in the previous structures of Orai.

**Figure 1.**
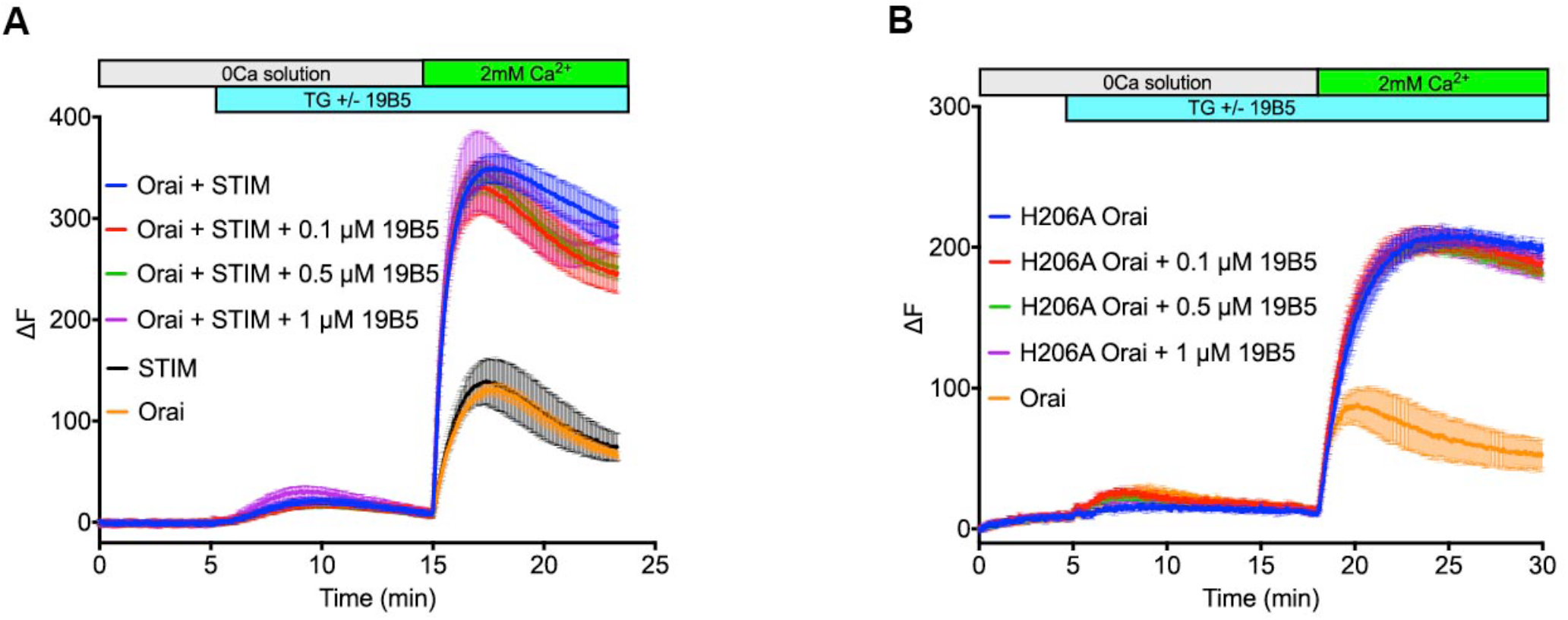
Analysis of the 19B5 antibody using a cellular Ca^2+^ influx assay. Ca^2+^ influx measurements were made from mammalian (HEK293) cells co-expressing wild type *Drosophila* Orai with *Drosophila* STIM (**a**) or expressing the H206A mutant of Orai alone (**b**). Cytosolic [Ca^2+^] levels were detected using a genetically encoded fluorescent Ca^2+^ indicator, GCaMP6s (Chen et al., 2013), as described previously (X. Hou, Burstein, & Long, 2018). Data are plotted as the change in fluorescence intensity (ΔF). Indicted concentrations of 19B5 antibody were used. Thapsigargin (TG), 2 mM CaCl_2_, and purified 19B5 antibody were added at the indicated times (horizontal bars). Controls (Orai or STIM alone) indicate background fluorescence levels. Standard error, derived from three independent measurements, is shown for each condition.

Having identified 19B5 as a candidate for structural analysis, we sought to investigate the effect of the antibody on the function of the channel. To do so we performed Ca^2+^ influx measurements in mammalian cells that expressed Orai_H206A_ or wild-type Orai and STIM components. The addition of 19B5 antibody had no discernable effect on Ca^2+^ influx through the channels (Figure 1 and Figure 1 - figure supplement 1). Therefore even though 19B5 binds to the extracellular side of the channel, it does not inhibit Ca^2+^ flux through the constitutively active Orai_H206A_ channel or through wild type Orai when activated by STIM.

### Cryo-EM structure determination of the Orai_H206A_ – Fab complex

In order to determine the structure of the Orai_H206A_ – Fab complex, cryo-EM data were collected from a sample of purified Orai_H206A_ that had been reconstituted into amphipols and combined with an excess of the 19B5 Fab antibody fragment (Methods). Classification of the particle images revealed Orai-Fab complexes containing one, two, or three Fabs bound per channel and some free Fab fragments (Figure 2 – figure supplements 1 and 2). We generated 3D reconstructions from particles that contained channels and found that these reconstructions contained six Orai subunits regardless of the number of Fab molecules that were bound, which indicted that the channels were uniformly composed of six Orai subunits in the sample (Figure 2 - figure supplement 2). The Fab molecules bound to the same epitope regardless of their stoichiometry with the channel (Figure 2 - figure supplement 2). As expected from biochemical analysis, this epitope was located on the extracellular side of the channel (Figure 2).

**Figure 2.**
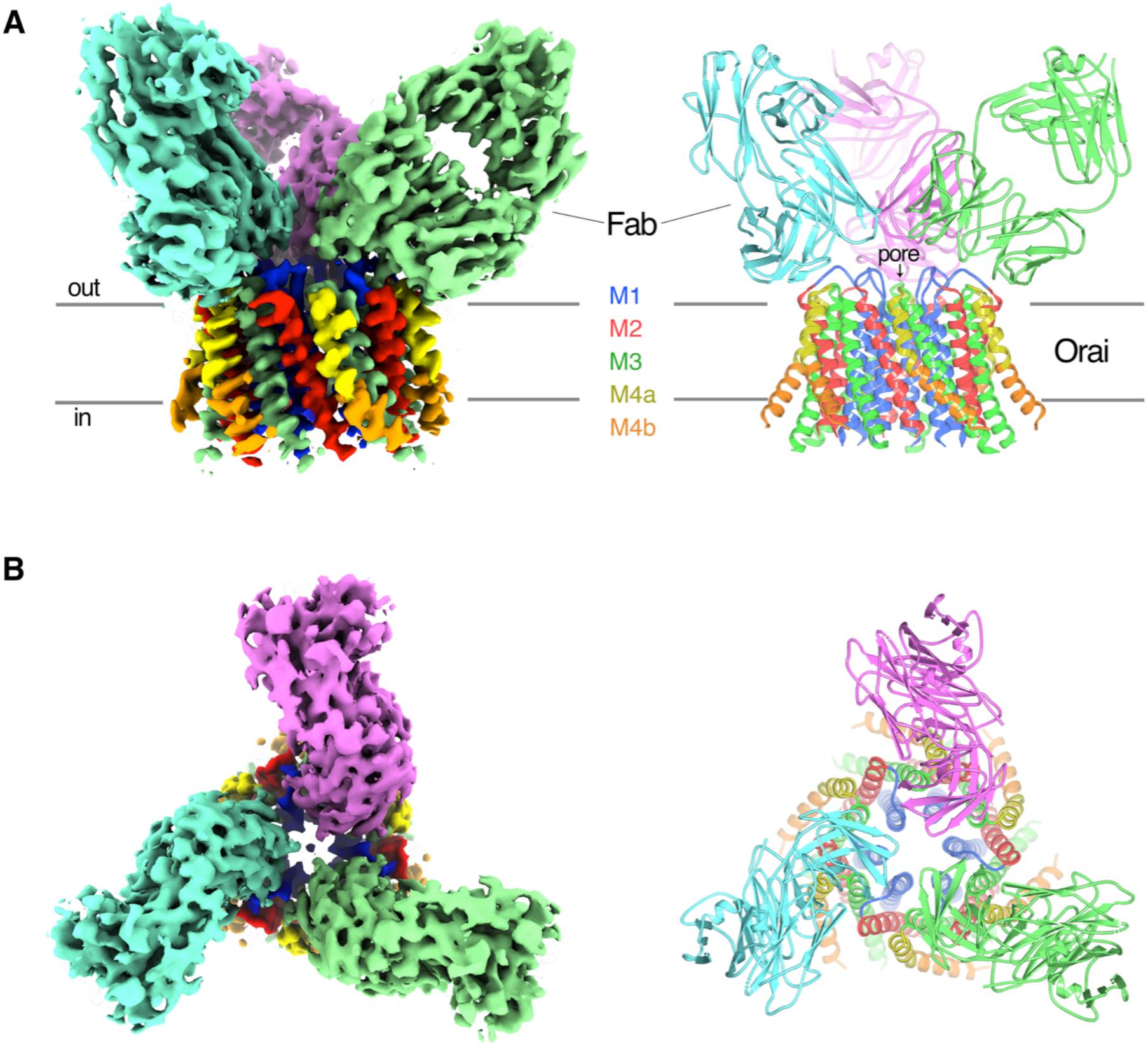
Orai_H206A_-Fab complex. **a, b**, Overall structure of the complex is shown from the perspective of the membrane (**a**) and from the extracellular side (**b**). The cryo-EM map (left) and the cartoon representation of the structure (right) are shown. Each Fab is colored individually; The α-helices of Orai are colored as indicated. Horizontal bars in (a) denote approximate boundaries of the plasma membrane.

Particles of Orai-Fab complexes that contained three Fabs yielded higher-resolution cryo-EM maps and were used for structure determination. On account of the antibodies, cryo-EM reconstructions displayed three-fold (C3) symmetry even when symmetry was not imposed (Figure 2 – figure supplement 1 and 2) and therefore C3 symmetry was incorporated during subsequent cryo-EM processing. The final 3D reconstruction is determined to 3.3 Å overall resolution (Figure 2, Figure 2 – figure supplements 1 and 3). Local resolution estimates and visual inspection of the maps indicate that the central region of the complex comprising the M1, M2, and M3 helices and the variable domains of the Fabs have the most well-defined density (at ∼ 3.1 Å resolution), while the M4 helices and the constant domains of the Fabs on the periphery are less well defined (Figure 2 - figure supplement 3). The atomic model has good stereochemistry and good correlation with the cryo-EM density (Figure 2 - figure supplements 3, 5). It contains the variable domains of three Fab molecules and amino acids 156-305 of Orai (except for the disordered M2-M3 loop, amino acids 217-239). The M4-ext helices (amino acids 306-341) and the N-terminal ends of the M1 helices (amino acids 144-156), which were observed in the X-ray structures, are not visible in the cryo-EM map, possibly due to flexibility in these regions and/or their location at the fringe of the cryo-EM reconstruction.

Each Fab binds to the extracellular side of Orai, near the entrance of the pore (Figure 2, Figure 2 - figure supplement 4). There are six identical binding sites for the Fabs, on account of the hexameric architecture of Orai. While there are no visible contacts between the antibodies, the channel can only accommodate up to three Fabs because of steric restraints (Figure 2). In accord with the Ca^2+^ influx results (Figure 1), there is adequate room for Ca^2+^ entry with Fabs bound (Figure 2). Each antibody primarily interacts with the connection between the M1 and M2 helices (Figure 2 - figure supplement 4). This connection, which was not visible in previous X-ray structures of Orai and comprises amino acids 179-189, forms a structured “turret” (Figure 3). Six turrets, from the six subunits, constitute the extracellular entrance of the pore (as described below). Each Fab predominately interacts with the M1-M2 turret of one subunit, but some contacts are also made with the neighboring Orai subunit (Figure 2, Figure 2 - figure supplement 4). These two adjacent turrets adopt indistinguishable conformations (Figure 2 - figure supplement 3). This correspondence, in spite of the different interactions between the Fab and the two turrets, suggests that the antibody has minimal effects on the structure of the turret. The property that the antibody preferentially binds to the intact channel rather than to denatured protein is an additional indication that the turret is a structured element of the channel. Weak density for the turret is present in our previous X-ray structures of Orai in both closed and opened conformations (Figure 4), and although this density was not strong enough to direct model building (X. Hou et al., 2018; Xiaowei Hou et al., 2012), the density is further evidence that the turrets are a structural component of the channel.

**Figure 3.**
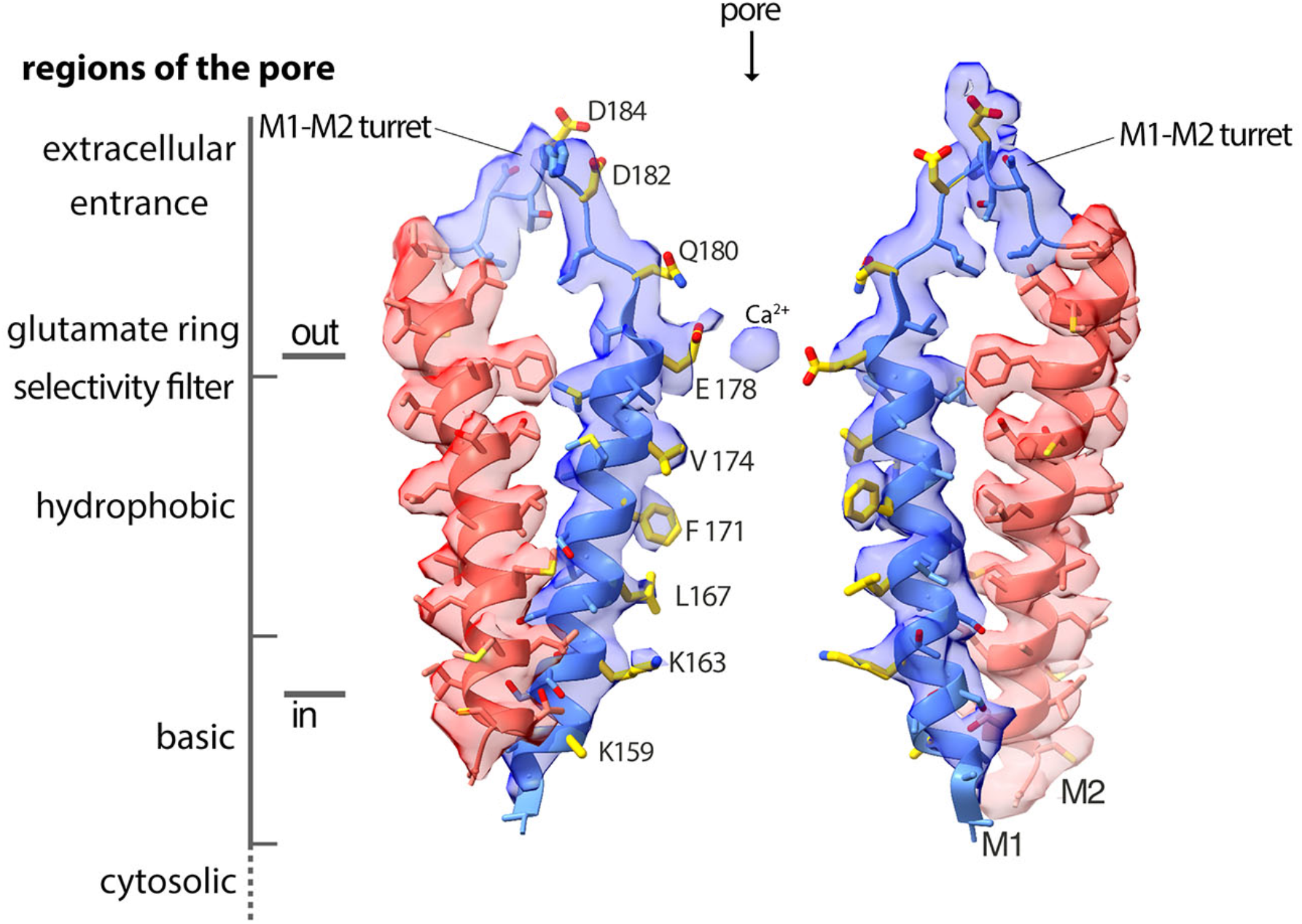
Cryo-EM density and structure of the pore. Cryo-EM density is displayed as semitransparent surface, showing the M1 through M2 portions of the channel from two opposing subunits (M3-M4 and other subunits are omitted for clarity). The atomic model is shown in cartoon representation, with amino acid side chains drawn as sticks. Amino acid side chains on M1 and on the M1-M2 turret that face the ion conduction pathway have yellow colored carbon atoms. Nitrogen and oxygen atoms are colored dark blue and red, respectively. Regions of the pore are indicated.

**Figure 4.**
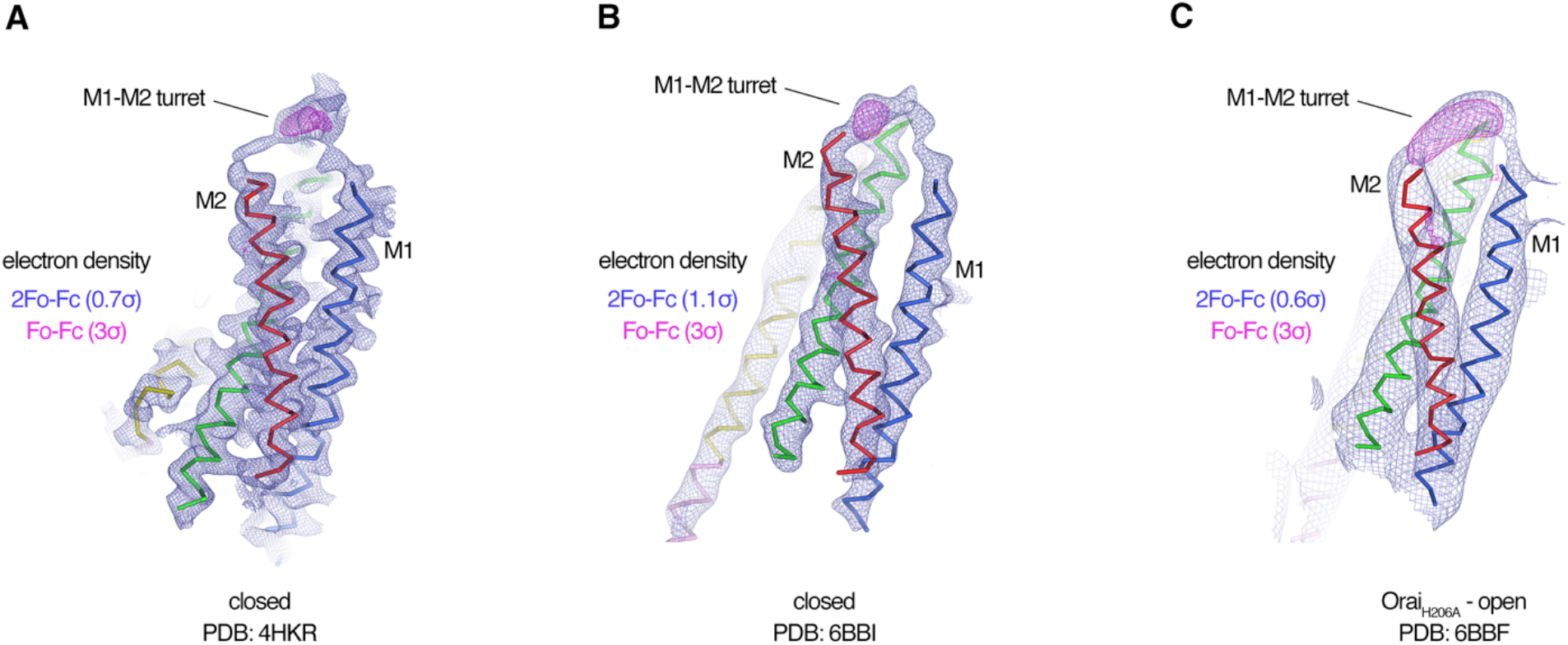
Evidence for the M1-M2 turret in previous X-ray structures of Orai. **a-c**, An Orai subunit (Cα representation) is shown from three X-ray structures. In two of the structures (a,b), the pore of Orai is closed (X. Hou et al., 2018; Xiaowei Hou, Pedi, Diver, & Long, 2012). The depiction in (c) is of the low-resolution X-ray structure of Orai_H206A_ (X. Hou et al., 2018). 2Fo-Fc and Fo-Fc density maps are drawn as blue and magenta mesh, respectively, at the indicated σ levels. The electron density maps were calculated using the map coefficients FWT/PHWT and DELFWT/DELPHWT, respectively, that can be downloaded from the RCSB Protein Data Bank using PDB IDs: 4HKR, 6BBI, and 6BBF, respectively. In (c), 24 fold averaging was applied to the map in real space according to the non-crystallographic symmetry present in this crystal form to increase the signal-to-noise level. Weak 2Fo-Fc density for the M1-M2 turret is visible in each of the three structures (at 0.7σ, 1.1σ, and 0.6σ, respectively). Positive densities in the Fo-Fc difference maps at 3σ confirm the presence of the M1-M2 turrets even though they were not built in the models due to relatively weak density and/or low resolution X-ray data. The structures in (a and b) are of Orai in a quiescent conformation and of K163W Orai in an unlatched-closed conformation, respectively. In those structures, the pores adopt indistinguishable closed conformations (the M4-ext helices have different conformations) (X. Hou et al., 2018; Xiaowei Hou et al., 2012).

The overall structure of the channel agrees with the low-resolution X-ray structure of Orai_H206A_ within the error of that structure (Figure 5a). This correspondence of 3D structures determined using different methodologies and under different conditions (detergent micelles were used for the X-ray structure) and the abilities of the antibody to bind both wild-type and H206A Orai are further indications that the antibody binds to a native structure of the channel.

**Figure 5.**
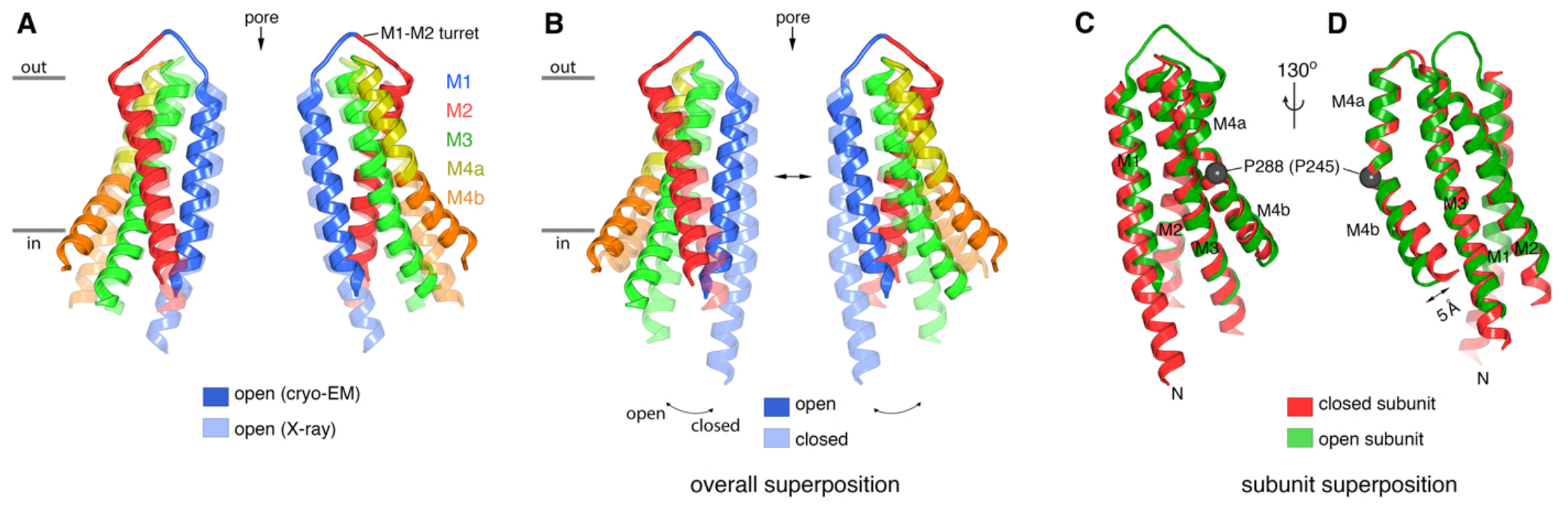
Comparison of the cryo-EM structure of Orai_H206A_ with other structures of the channel. **a**, Comparison between the cryo-EM structure and a previous low-resolution X-ray structure of Orai_H206A_ (X. Hou et al., 2018). Two opposing subunits of Orai are depicted in cartoon representation and colored as indicated (four subunits are omitted for clarity). The X-ray structure (PDB: 6BBF) is colored in lighter shades. The M1-M2 turrets revealed by the cryo-EM structure, which create the extracellular entrance of the pore, are indicated. The mainchain RMSD between the structures is 1.5 Å. **b**, Comparison between the cryo-EM structure of Orai_H206A_ and a 3.3 Å resolution X-ray structure of a closed conformation (quiescent conformation, PDB: 4HKR). The mainchain RMSD between these two structures is 2.7 Å. The depiction is as in (a), with the closed structure drawn in lighter shades. Arrows highlight dilation of the pore and the tilting of subunits. Superpositions in (a) and (b) were made by aligning complete hexameric channels. **c, d**, Superposition of an individual subunit from Orai_H206A_ with an individual subunit of the closed structure (PDB: 4HKR), shown in two orientations (c, d). The mainchain RMSD between these two subunits is 0.74 Å. Aside from a slight displacement of the C-terminal portion of M4b (arrow), the conformations of an isolated subunit are highly similar between the open and closed structures.

### The extracellular entrance

The six M1-M2 turrets, one from each subunit, constitute the extracellular entrance of the pore (Figures 3, 5). The amino acid sequence of the M1-M2 turret and its length are highly conserved among Orai channels (Figure 6). The cryo-EM structure establishes that the M2 helix of a given subunit is located directly behind the M1 helix of the same subunit (Figures 3, 5a). The polypeptide of the turret adopts an extended secondary structure that connects the C-terminal end of M1 (at Glu 178) with the N-terminal end of M2 (at Gly 190). Each turret extends 20 Å above Glu 178 and has a shape somewhat like that of an inverted “V”. The cryo-EM structure allows us to visualize that the C-terminal end of the M1 α-helix is stabilized by hydrogen bonds between backbone oxygen atoms (at Met 176 and Val 177) and Lys 270 from the M3 helix of a neighboring subunit (Figure 6d). The amino acid sequence conservation of Lys 270 and the region surrounding it suggests that the corresponding residue in human Orai1, Lys 178, participates in similar interactions (Figure 6e).

**Figure 6.**
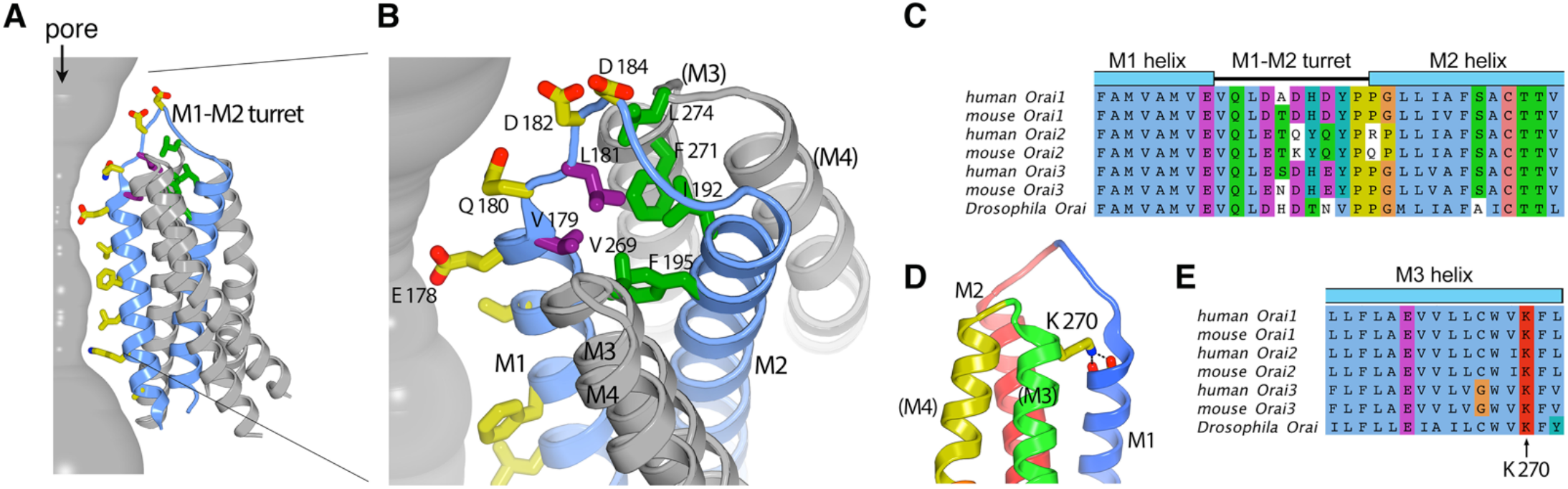
The M1-M2 turret. **a, b**, Overall and close-up views highlighting the M1-M2 turret. The M1 and M2 helices and the intervening turret from a single subunit are light blue. The M3 and M4 helices from the corresponding subunit and a neighboring one are colored gray, with parentheses indicating helices from the neighboring subunit. Select residues are drawn as sticks: pore-lining amino acids (yellow carbon atoms), hydrophobic residues on the turret (purple), and cluster of hydrophobic residues beneath the turret (green). A section of the pore is depicted as a gray surface. **c**, Sequence alignment in the turret region. Clustal coloring; portions of the M1 and M2 helices nearest the turret are shown. **d**, Close up view showing an interaction between Lys 270 and the C-terminal end of M1. Lys 270 is drawn as sticks; hydrogen bonds with the backbone carbonyl oxygen atoms at the C-terminal end of M1 are depicted as dashed lines. Parentheses indicate that the M3 and M4 helices are from a neighboring subunit. **e**, Sequence alignment near Lys 270.

The N-terminal end of the turret contains a conserved “VQLD” motif (Val 179, Gln 180, Leu 181, and Asp 182) that is located immediately adjacent to Glu 178 of the selectivity filter (Figure 6a-c). The two hydrophilic residues of the motif, Gln 180 and Asp 182, are oriented toward the pore and the aqueous environment of the extracellular entrance. The two hydrophobic amino acids, Val 179 and Leu 181, point away from the aqueous environment and participate in a network of hydrophobic interactions that appear to provide structural stability to the turret (Figure 6a,b).

The extracellular entrance of the pore created by the turrets is considerably larger than was appreciated from the previous structures of Orai (Figure 7). The entrance is funnel shaped, with a diameter of ∼20 Å at its widest point, and narrows to join with the selectivity filter of the channel at Glu 178. Gln 180, Asp 182 and Asp 184 contribute to the walls of the funnel and together with Glu 178 produce an electrostatic surface that is exceedingly negative on the extracellular side and would tend to increase the local concentration of cations within it (Figure 8).

**Figure 7.**
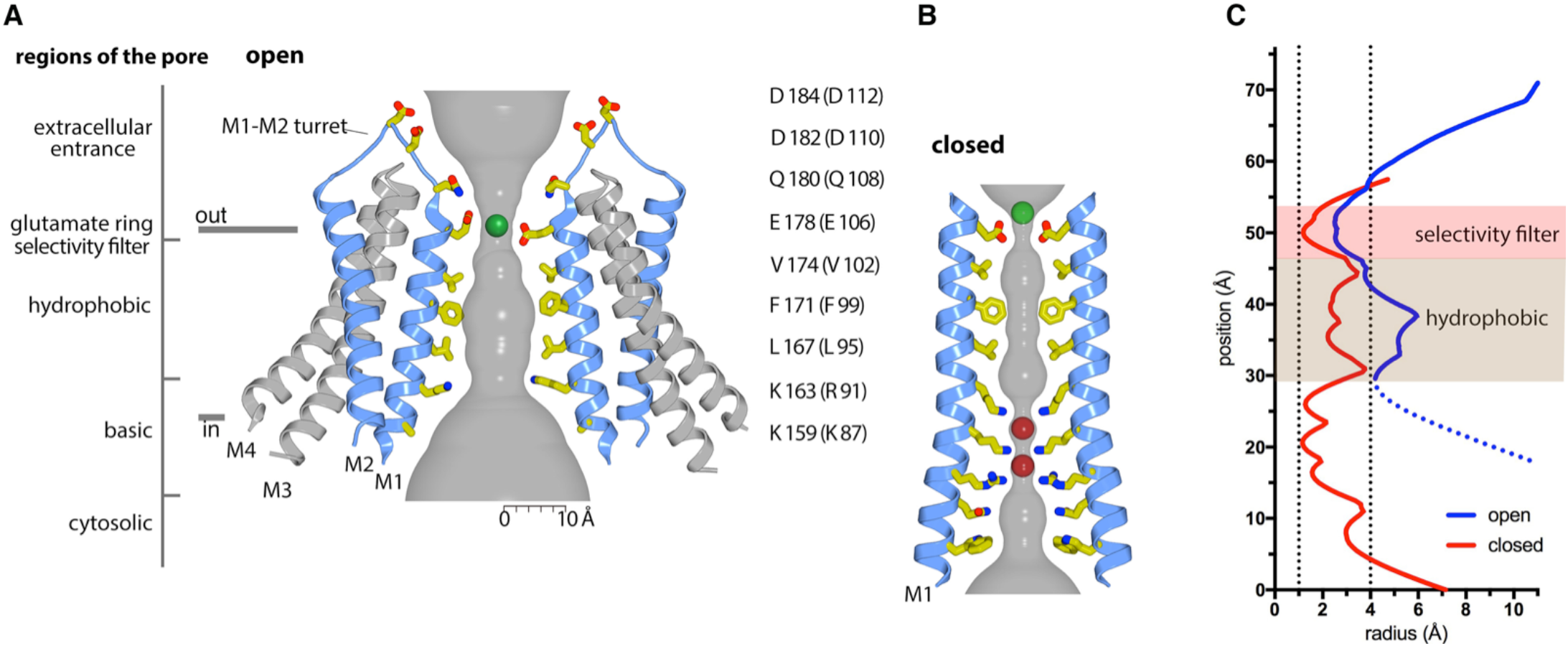
Ion pore and gating transitions. **a**, Open pore of Orai_H206A_. Two opposing subunits are shown as cartoons around the ion pore, which is depicted as a gray surface and represents the minimal radial distance to the nearest van der Waals contact. Amino acids lining the pore are shown as sticks (yellow carbon, red oxygen, and blue nitrogen atoms). These amino acids are labeled (the side chain of K159 is only partially modeled due to weak density); parentheses denote human Orai1 counterparts. A green sphere indicates Ca^2+^. **b**, Closed pore of Orai. Two opposing M1 helices from the X-ray structure of the quiescent conformation (PDB: 4HKR) are shown (ribbons) with amino acids that line the pore drawn as sticks. The ion pore is depicted as in (a); red spheres represent iron/anion binding sites. **c**, Comparison of the dimensions of the closed and open pores. Pore radius (x-axis) denotes the minimal radial distance to the nearest van der Waals contact along the axis of the pore (y-axis). The positions and scale along the y-axis correspond to (a,b). The selectivity filter and hydrophobic regions are shaded. The dotted line of the open trace represents the basic region, in which there is more uncertainty due to weaker cryo-EM density. The ionic radius for dehydrated Ca^2+^ (1.0 Å) and the radius of a hydrated Ca^2+^ ion (approximately 4 Å) are indicated as vertical dashed lines.

**Figure 8.**
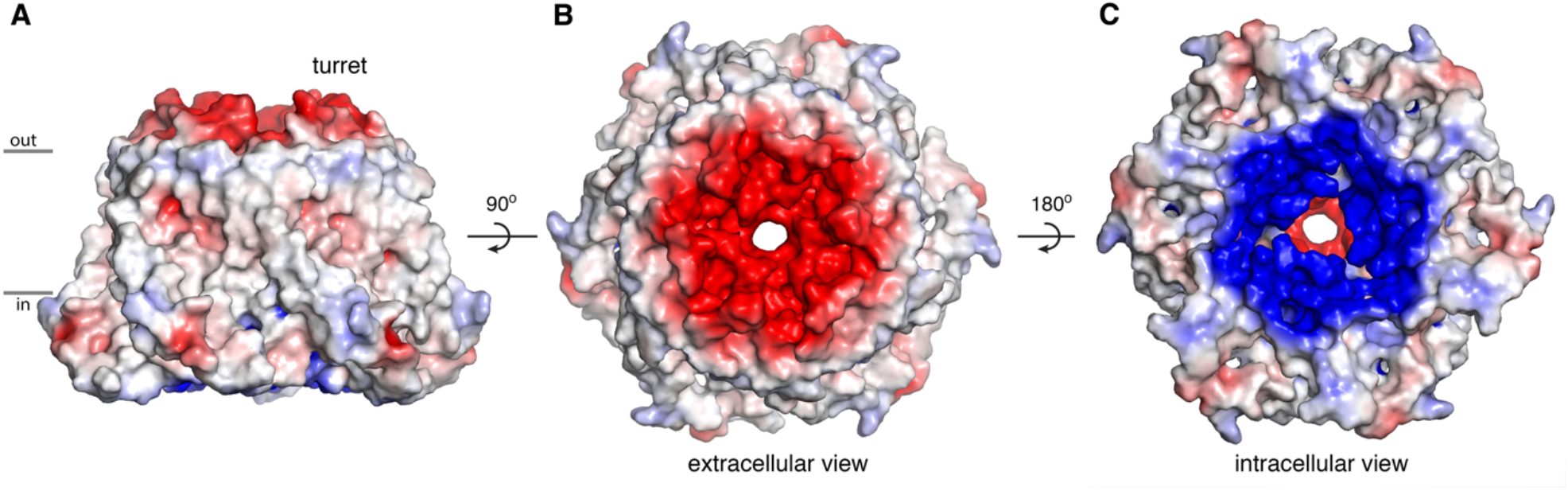
Electrostatic surface. **a, b, c**, Molecular surface of Orai_H206A_ viewed from the membrane (a), extracellular (b) and intracellular (c) perspectives. Coloring is according to the electrostatic potential, which is contoured from –5 kT (red) to +5 kT (blue) (dielectric constant: 80).

### The open pore

With the additional appreciation of the extracellular entrance provided by the cryo-EM structure, the pore has five distinct sections (Figure 3 and 7a). From the extracellular to the intracellular side, these are: the extracellular entrance composed of the M1-M2 turrets, the selectivity filter, a hydrophobic section, a basic section, and a cytosolic section. Aside from residues on the turret, the walls of the pore are formed by the M1 helices. As such, the amino acid side chains emanating from M1 create the physiochemical environment for the majority of the pore. Most of these pore-lining side chains are resolved in the cryo-EM map, providing a detailed view of the open pore of Orai_H206A_ (Figure 3). Figure 7a shows the approximate dimensions of the open pore and indicates the residues lining its walls. Changes are evident along the length of M1 in comparison to the closed conformation of the pore (Figure 7b). These changes increase the diameter of the pore along its entire length.

A “glutamate ring” of Glu 178 residues from the six subunits is the narrowest constriction of the open pore (Figure 7a,c). Previous functional and structural data indicate that the glutamate ring forms the selectivity filter of the channel that is responsible for the channel’s high-selectivity for Ca^2+^ (Xiaowei Hou et al., 2012; Prakriya et al., 2006; Vig et al., 2006; Yeromin et al., 2006). Cryo-EM densities for the side chains of the Glu 178 residues are relatively weak in comparison to other amino acids on the M1 helices, as is often the case for negatively charged amino acids in cryo-EM structures owing to interactions with the electron beam (Figure 3, Figure 2 - figure supplement 3a). Nevertheless, the densities suggest that the side chains of the Glu 178 residues adopt two favorable rotamer conformations, with one glutamate residue in an “up” rotamer and an adjacent glutamate in a “down” rotamer, such that the amino acids alternate in an up-down manner around the hexameric pore (Figure 3, Figure 2 - figure supplement 3). Density consistent with Ca^2+^ is present at the center of the glutamate ring (Figure 3). The density corresponds to the previously identified binding location of Ba^2+^, a permeable surrogate for Ca^2+^, that was identified from X-ray studies of Orai_H206A_ (X. Hou et al., 2018).

In the closed conformation of the pore observed in the X-ray structure of wild-type Orai, all six of the Glu 178 residues are modeled in a “down” rotamer conformation (Figure 7b) (Xiaowei Hou et al., 2012). It should be noted, however, that there is a degree of ambiguity regarding the precise conformations of these residues in both the X-ray and cryo-EM structures. It is possible that the side chains are dynamic and do not adopt a particular conformation for an extended period of time. For example, there are no additional hydrogen bonds between the glutamate residues and other amino acids that might stabilize their conformations. This is in marked contrast to another highly selective Ca^2+^ channel, the mitochondrial Ca^2+^ uniporter, in which the conformations of a ring of glutamate residues in its selectivity filter are stabilized by van der Waals interactions and hydrogen bonds with other amino acids (Baradaran, Wang, Siliciano, & Long, 2018; C. Fan et al., 2018; M. Fan et al., 2020; Nguyen et al., 2018; C. Wang, Jacewicz, Delgado, Baradaran, & Long, 2020; Y. Wang et al., 2019; Yoo et al., 2018). In spite of some ambiguity regarding the conformations of the glutamate side chains in the selectivity filter of Orai, the helical backbone of the M1 helix at the glutamate ring is shifted away from the central axis of the pore by approximately 1 Å relative to the closed conformation (Figure 5b, Figure 7). Thus, opening of the channel involves a degree of dilation at the selectivity filter (Figure 7).

Three perfectly conserved residues that are positioned on successive helical turns of each M1 helix, Leu 167, Phe 171, and Val 174 (Leu 95, Phe 99, and Val 102 in human Orai1), comprise a hydrophobic region of the pore (Figure 7). The entire hydrophobic region is markedly wider in the open conformation than when the pore is closed: relative to the closed conformation the helical backbone of each M1 helix is displaced by more than 2 Å away from the center of the pore within it (Figure 7).

Changes within the hydrophobic region at and around Phe 171 are particularly noteworthy. In the closed conformation, the side chains of the six Phe 171 residues (from the six subunits) are in close proximity and are located near the central axis of the pore (Figure 9b, Figure 9 – figure supplement 1). Their conformations are stabilized by hydrophobic packing among themselves and by close van der Waals contacts with the Gly 170 residues of neighboring subunits (Figure 9b). The Cα position of the perfectly conserved Gly 170 residue (Gly 98 in Orai1) is not exposed to pore in the closed state (Figure 9b). Opening increases the exposure of Gly 170 to the pore by an outward sliding movement of Phe 171 from the adjacent subunit that results from outward movements of the subunits (Figure 9e). Other than the increased exposure of Gly 170 to the pore in the open state, all of the amino acids that line the walls of the pore in the closed conformation also do so in the open conformation. Notably, the opening transition does not involve appreciable rotation of the M1 helices. Rather, rigid body movement of the helices away from the center of the pore constitutes opening (Figure 5b,c, Figure 7, Figure 9, Videos 1 and 2).

**Figure 9.**
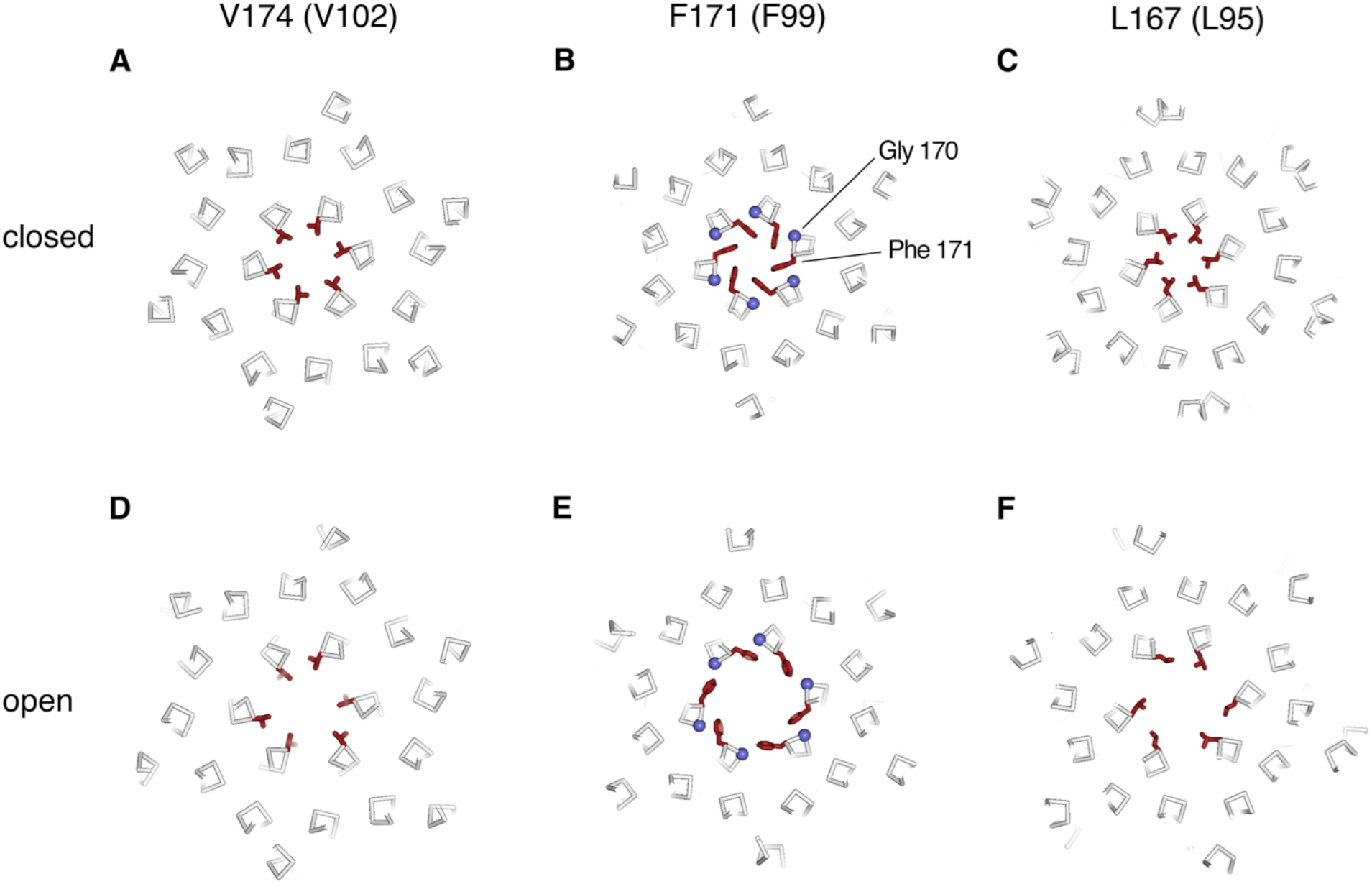
Conformational changes in the hydrophobic region. **a**-**f**, Slices through the hydrophobic region of the pore from the closed (PDB 4HKR) and Orai_H206A_ open structures (upper and lower panels, respectively). The structures are represented as ribbons with amino acids side chains in the hydrophobic region drawn as sticks (dark red). Gly 170 is depicted as a blue sphere. The perspectives are from the extracellular side, perpendicular to the pore, which is centrally located. The slices correspond to approximately 4 Å slabs centered at Val 174, Phe 171, and Leu 167, as indicated. Inspection of the upper and lower panels indicates the outward rigid body movements of subunits. Videos 1 and 2 depict these conformational changes.

Oddly for a cation channel, the pore of Orai contains an extremely basic region, which is composed of eighteen lysine or arginine residues (three residues from each of the six subunits) and located just below the hydrophobic region (Figure 7). These amino acids (Arg 155, Lys 159, and Lys 163 in *D*. Orai) are conserved as lysine or arginine residues in all Orai channels (the corresponding amino acids are Arg 83, Lys 87, Arg 91 in human Orai1). In the closed state, these amino acids are in close proximity and form binding sites for anions (Xiaowei Hou et al., 2012). Although the identity of the physiological anion is not yet established, X-ray and mass-spectrometry analysis indicate that purified wild type Orai contains an iron complex within the anion binding site that may be (FeCl_6_)^-3^ (Xiaowei Hou et al., 2012). The basic region is markedly wider in the open conformation owing to the outward movement of the M1 helices; the Cα positions of Lys 159 on opposing M1 helices are 12 Å further apart than in the closed structure (Figure 7). As was the case in the X-ray structure of Orai_H206A_ (X. Hou et al., 2018), density for the anion/iron complex is not present within the widened basic region in the cryo-EM structure. Although side chain densities for the basic residues are less well-defined than for other residues on M1, which suggests their flexibility, the helical register of M1 and some side-chain density for Lys 163 (Figure 3) indicate that the basic residues are exposed to the pore in the open state. We hypothesize that small cellular anions such as chloride would shield the basic amino acids from Ca^2+^ ions permeating through the open channel.

### Gating conformational changes in the channel

The changes between the closed and open conformations of the channel are best described as rigid-body shifts in which the M1-M4 portion of each of the six subunits moves away from the center of the pore (Figures 5, 7 and 9, and Videos 1 and 2). For instance, an ∼ 2.5 Å outward movement of the M1 helix at Phe 171 within the hydrophobic region is accompanied by outward movements of similar magnitude for the M2, M3 and M4 helices (∼ 1.6 Å, ∼ 2.0 Å and ∼ 1.7 Å measured at residues within the same horizontal plain as Phe 171, respectively) (Figure 9b,e). The outward movement is more dramatic on the cytosolic side of the channel and this results in slight tilting of subunits away from the pore on that side (Figure 5b, Video 2). In addition to these rigid body motions, a bend in the M4 helix at Pro 288, which delineates it into M4a and M4b, is less bent in the cryo-EM structure of Orai_H206A_ than it is in the quiescent conformation. This results in ∼5 Å outward movement of each of the M4b helices from its position in the quiescent conformation (measured at Ser 303) (Figure 5b,d).

His 206, the amino acid that when mutated to alanine gives rise to the constitutively activated channel, is located on M2 and would not contribute to the walls of the ion pore in either the closed or the open conformation (Figure 9 - figure supplement 2). In the WT channel, His 206 makes a hydrogen bond with Ser 165 from M1 and is within a network of van der Waals interactions that involves residues from M1, M2 and M3 (Irene Frischauf et al., 2017; X. Hou et al., 2018; Xiaowei Hou et al., 2012; Yeung et al., 2018). The H206A mutation eliminates this hydrogen bond and diminishes the van der Waals interactions (Figure 9 - figure supplement 2c), and this evidently alters the free energy profile of the channel sufficiently to favor an open state in the absence of STIM. A histidine can be readily modeled back into the Orai_H206A_ structure, which suggests that the wild type channel would be able to adopt the observed conformation (e.g. when activated by STIM) (Figure 9 - figure supplement 2b).

Remarkably, the opening observed in Orai_H206A_ structure does not involve notable side chain conformational changes. This can be appreciated by comparing amino acid conformations in Figure 9 and the opening transition depicted in Videos 1 and 2. Because the helices of Orai do not interdigitate and are nearly perpendicular to the membrane, we predict that the movement of an individual subunit would be relatively independent of neighboring subunits.

## Discussion

The cryo-EM structure of Orai_H206A_ provides near-atomic detail of an open conformation of the channel. Somewhat surprisingly, opening of the pore does not involve twisting or bending movements of transmembrane helices that would be analogous to gating movements of many other cation channels such as voltage-dependent potassium, sodium or calcium channels as was first exemplified by structural studies of potassium channels (Jiang et al., 2002b; MacKinnon, 2003), nor does it involve marked rotation along helical axes. Rather, opening observed in Orai_H206A_ involves rigid-body outward movement of each subunit. The amino acids within the transmembrane region adopt similar conformations in the closed and the open conformations (Video 1). The outward movement of the subunits results in a dilation of the pore and repositioning of Phe 171, which slides away from the central ion pathway to widen the hydrophobic region of the pore (Figure 9 and Videos 1 and 2).

The open structure of the pore is in remarkable agreement with the amino acids predicted to contribute to the pore from cysteine accessibility experiments (McNally, Yamashita, Engh, & Prakriya, 2009; Megumi Yamashita et al., 2017; Yubin Zhou, Ramachandran, Oh-hora, Rao, & Hogan, 2010). In spite of the potential disruptive effects caused by the introduction of cysteine mutations within a tightly packed and highly conserved ion pore, all of the residues that were predicted to contribute to the pore from these experiments (Arg 91, Leu 95, Gly 98, Phe 99, Val 102, Glu 106 of human Orai1) are observed to do so in the structures. The accessibility profile of a cysteine substituted for Gly 98 (Gly 170 in *D*. Orai) was particularly intriguing. In the background of the pore-lining V102A mutation, which is constitutively active (McNally, Somasundaram, Yamashita, & Prakriya, 2012) but hypothesized to be structurally similar to the closed conformation of the channel (Megumi Yamashita et al., 2017), channels bearing a cysteine substitution of Gly 98 were not blocked when Cd^2+^ was applied to the extracellular side, but they were blocked when the mutant channel was activated by STIM (Megumi Yamashita et al., 2017). To account for this change in accessibility to Cd^2+^, it was hypothesized that the M1 helix rotates modestly (by ∼ 20°) along its helical axis when the channel opens (Megumi Yamashita et al., 2017). We do not observe a marked rotation of M1 between the closed and open structures, but we do observe a change in the exposure of the corresponding residue, Gly 170 (Gly 98 in Orai1), that could explain the observed accessibility profile. In the closed state, Gly 170 is hidden behind Phe 171 and is therefore not exposed to the pore (Figure 9, Video 1). However, Gly 170 becomes exposed to the pore in the open conformation due to the outward movements of the subunits and the sliding motion of Phe 171 relative to it (Figure 9 and Videos 1,2). Yamashita et al. observed a change in the ability of Cd^2+^ to block the F99C mutant of Orai1 - the cysteine substitution of Phe 99 in Orai1 (corresponding to Phe 171 in Orai) was blocked by Cd^2+^ in the V102A closed state but was markedly less blocked in the STIM-activated state (Megumi Yamashita et al., 2017). This finding at first seems contradictory to the structure because Phe 171 is exposed to the pore in both the open and closed 3D structures. However, a possible explanation for the accessibility profile of the F171C mutation is that the cysteine residues on opposing sides of the pore would be too far apart in the open conformation of the pore (∼ 17 Å measured between sulfur atoms) to be coordinated by Cd^2+^, which typically has a S-Cd^2+^ coordination distance of ∼ 2.5 Å and requires multiple sulfur ligands for efficient binding (Rulisek & Vondrasek, 1998). In general agreement with the structural observations, molecular dynamics simulations of the H206A mutation have suggested that the hydrophobic region of the pore widens when the channel opens (Irene Frischauf et al., 2017; M. Yamashita et al., 2020; Megumi Yamashita et al., 2017; Yeung et al., 2018). Although the details of the widening process (e.g. rotation vs. dilation) are slightly different between the cryo-EM structure and molecular dynamics simulations, the structure of Orai_H206A_ supports the growing consensus that widening of the hydrophobic region underlies gating and it provides a structural basis for its mechanism (as discussed in more detail below).

Liu et al. recently described low resolution X-ray and cryo-EM structures of *D*. Orai (determined at resolutions of ∼ 4.5 Å and ∼ 5.7 Å, respectively) with the mutation P288L (Liu et al., 2019), which has an activating phenotype in cells (Liu et al., 2019; Nesin et al., 2014; Palty, Stanley, & Isacoff, 2015). The authors propose that their structures reveal an open conformation of the pore (Liu et al., 2019), but we suspect that the pore is actually closed (non-conductive) in these structures. Side chain conformations are not visible owing to the low resolutions of theses structures, but the densities for α-helices from both the cryo-EM and X-ray structures of P288L Orai reveal a conformation that is highly similar to an “unlatched-closed” conformation that we observed previously (X. Hou et al., 2018) in which the pore is closed (Figure 9 - figure supplement 3b,c). Most notably, the hydrophobic region is narrow in the P288L structure as it is in the structures of Orai with a closed pore (X. Hou et al., 2018; Xiaowei Hou et al., 2012). Density consistent with an anion plug, which is a structural hallmark of the closed pore (X. Hou et al., 2018; Xiaowei Hou et al., 2012), is also observed in the P288L structures (Figure 9 - figure supplement 3c). While the corresponding P245L mutation of human Orai1 causes activation of the channel when expressed in mammalian cells (Liu et al., 2019; Nesin et al., 2014; Palty et al., 2015), we have not been able to detect ion permeation through purified P288L Orai in liposomes, which suggests that the channel is not constitutively open on its own (Figure 9 - figure supplement 3a). We suspect that channel activation by this mutation in cells may be dependent on endogenous STIM molecules. Another gain-of-function mutation of human Orai1, T184M, has been reported to require STIM for its activation (Bohm et al., 2017). Further analysis of these intriguing mutations may shed light on the coupling between STIM and Orai and the mechanisms for how STIM activates the channel following Ca^2+^ store depletion.

The M1-M2 turrets observed in the cryo-EM structure of Orai_H206A_ reveal that the channel has an electronegative extracellular entrance that is wide enough to accommodate hydrated ions. The negatively charged residues on the walls of the entrance (Asp 182 and Asp 184, corresponding to Asp 110 and Asp 112 in human Orai1) would tend to concentrate Ca^2+^ and other cations within vicinity of the pore, as has been proposed on the basis of functional studies of mutations of these residues and molecular dynamics simulations (I. Frischauf et al., 2015). In agreement with the observed structural locations of Asp 182 and Asp 184, mutation of either residue to alanine reduces the affinity of Gd^3+^ block but has no detectable effect on Ca^2+^ selectivity, which is mostly conferred by Glu 178 (Prakriya et al., 2006; Vig et al., 2006; Yeromin et al., 2006). (Glu 178, Asp 182 and Asp 184 of *Drosophila* Orai are referred to as Glu 180, Asp 184 and Asp 186, respectively in (Yeromin et al., 2006)). Additionally, experiments using cysteine substitutions in the M1-M2 loop of Orai1 reveal a pattern of accessibility to thiol-binding compounds (McNally et al., 2009) that suggests that the M1-M2 connections in human Orai1 channels are structured similarly to the turrets in the Orai_H206A_ structure and form an analogous extracellular entrance. In those experiments, cysteine substitutions of the hydrophobic residues corresponding to Val 179 and Leu 181 in Orai (Val 107 and Leu 109 in Orai1), which are mostly buried in the structure, underwent modification by the cysteine-reactive compound MTSET (2- (trimethylammonium)ethyl methanethiosulfonate) approximately 10-fold more slowly than cysteine substitutions of the hydrophilic residues Gln 180 and Asp 182 (Gln 108 and Asp 110 in Orai1) that form the walls of the extracellular entrance in the structure. This cysteine accessibility profile suggests that the hydrophilic residues Gln 108 and Asp 110 of Orai1 are more exposed to the pore than the hydrophobic ones (Val 107 and Leu 109 in Orai1), which is in accord with what we observe in the structure. In further agreement with the structure, the authors of that study concluded that the M1-M2 loops “flank a wide outer vestibule” of the channel (McNally et al., 2009).

The cryo-EM structure shows that the selectivity filter widens by 1-2 Å when the pore opens (Figure 7). The widening of the filter is in agreement with several lines of evidence. Mutation of Val 102 in human Orai1 (Val 174 in Orai) to serine, cysteine, alanine or other small amino acids creates channels that conduct cations without activation by STIM but which are considerably less selective for Ca^2+^ (McNally et al., 2012). Ion selectivity for Ca^2+^ is restored when STIM engages the mutant channel, which suggests that the opening of the channel by STIM induces a change in the selectivity filter (McNally et al., 2012). In additional evidence for a structural change in the filter, we have observed the repositioning of density for barium ions (Ba^2+^) within the filter between the X-ray structures of the closed channel and Orai_H206A_ (X. Hou et al., 2018; Xiaowei Hou et al., 2012). Ba^2+^, which is conductive in Orai, binds above the filter when the channel is closed, but seems to bind within the filter in the open pore. The high-resolution structures support this repositioning of Ca^2+^ in the filter (Figure 7). Finally, spectroscopic studies suggest that changes occur at the extracellular side of the pore when Orai opens (Gudlur et al., 2014). Thus, unlike classical potassium channels in which the selectivity-filter can adopt a discrete conformation (to first approximation) regardless of whether the activation gate is open or closed (Jiang et al., 2002a; Long, Tao, Campbell, & MacKinnon, 2007; Y Zhou, Morais-Cabral, Kaufman, & Mackinnon, 2001), the selectivity filter of Orai adopts a discernably different conformation in an open conformation of the channel. The cryo-EM structure does not provide further details regarding ion coordination or the number of Ca^2+^ ions that can be accommodated in the filter, owing to the relatively weak density for the Glu 178 side chains. However, we suspect that ion selectivity involves a classical ion-selectivity mechanism in which two Ca^2+^ ions are positioned in single-file within the filter (X. Hou et al., 2018; Sather & McCleskey, 2003). Flexibility of the glutamate side chains, with a fraction of them in an “up” rotamer and a fraction in a “down” rotamer, as is suggested by the cryo-EM structure, may facilitate accommodation of two Ca^2+^ ions. This hypothesis is consistent with the observation that channels composed of concatemers of alternating wild type human Orai1 subunits and subunits containing the E106Q mutation conduct cations but have reduced selectivity for Ca^2+^ (X. Cai et al., 2018), whereas introducing this mutation into every subunit yields a non-conductive channel (Prakriya et al., 2006; Vig et al., 2006).

We suspect that there is fluidity in the conformations of amino acids on the pore such that the pore environment is not perfectly symmetric at any given time. For instance, the selectivity filter allows mixed conformations of the Glu 178 side chains. The conformations of the basic region may be particularly variable among subunits as density for this region is weaker than for other regions of the channel. We suspect that the cytosolic region of M1, which is mostly disordered in the cryo-EM structure, may adopt a more defined conformation upon the binding of STIM, which is thought to interact with this region to help stabilize the open pore (Derler et al., 2013; McNally, Somasundaram, Jairaman, Yamashita, & Prakriya, 2013; H. Zheng et al., 2013).

The X-ray and cryo-EM structures of Orai_H206A_ reveal that the basic region of the pore is markedly wider in the open conformation and no longer contains the anion plug that is observed when the pore is closed (X. Hou et al., 2018; Xiaowei Hou et al., 2012). Dilation of the basic region is consistent with previous biochemical experiments and molecular dynamics simulations that detected a widening of the pore in both the hydrophobic and basic regions in the activating H134A mutant of human Orai1 that corresponds to the H206A mutation of *D*. Orai (Irene Frischauf et al., 2017). However, several lines of evidence suggest that the basic region does not act as the primary impediment to ion permeation in the closed conformation of the channel. Richard Lewis and colleagues have shown that Orai1 channels in which any of the basic residues has been replaced by an acidic amino acid still form CRAC channels that require STIM for activation (Mullins & Lewis, 2016; Mullins, Yen, & Lewis, 2016). Interestingly, such mutations eliminate Ca^2+^-dependent inactivation, which suggests a role for the basic region in that process (Mullins & Lewis, 2016; Mullins et al., 2016). We have shown that removal of the basic region by replacement of all three residues with serine (R155S, K159S, K163S mutations) does not generate a constitutively active channel (X. Hou et al., 2018). Recent work by Murali Prakriya and colleagues showed that these serine substitutions also prevent activation of the channel by STIM, in further support that the basic region is important for the ability of the channel to open and/or conduct Ca^2+^ but suggesting that the basic region does not form the primary gate of the channel (M. Yamashita et al., 2020). We have shown that I^-^ can bind in the opened basic region and presume that similar anions such as Cl^-^ can do so (X. Hou et al., 2018). We hypothesize that permeation of Ca^2+^ through the basic region involves charge shielding by cellular anions, as has also been proposed on the basis of molecular dynamics simulations (Dong, Fiorin, Carnevale, Treptow, & Klein, 2013).

From the structures of Orai in open and closed conformations and the aforementioned functional and molecular dynamics studies (reviewed in (Yeung et al., 2020)), there is a consensus that the hydrophobic region of the pore functions as the primary impediment to ion permeation that controls Ca^2+^ flow. In the closed conformation, the hydrophobic region is too narrow and too hydrophobic to permit Ca^2+^ permeation, whereas the widened hydrophobic region in the Orai_H206A_ structure would allow it. The hydrophobic nature of this region is particularly important to prevent ion conduction in the closed conformation: substitutions of hydrophobic amino acids within it with smaller or more hydrophilic ones (e.g. the V102A, F99C mutations of Orai1) allow cation permeation without activation by STIM (McNally et al., 2012; Megumi Yamashita et al., 2017). Molecular dynamics simulations of carbon-nanotube model systems and ion channels suggest that the ability of water molecules to be accommodated, even transiently, within hydrophobic regions is associated with their abilities to conduct ions (Aryal, Sansom, & Tucker, 2015; Jensen et al., 2010). These studies suggest that when the diameter of a hydrophobic region is less than a certain width, the region becomes “dewetted” in that water molecules are rarely observed in it, and ion permeation is prevented. Conversely, when the diameter is large enough, water molecules are accommodated and ion permeation is permitted. From simulations using a hydrophobic constriction similar in length to the hydrophobic section in the pore of Orai, a steep transition occurs at a diameter of approximately 9 Å (Aryal et al., 2015). When the diameter is 8 Å or less, the hydrophobic section is dewetted most of the time, whereas water molecules are observed within it at even slightly larger diameters. This corresponds well with the structures of Orai: the diameter of the hydrophobic region is 5-6 Å in the closed state and 9-10 Å in the H206A open state (Figure 7). The simulations also show that increasing the hydrophilicity of a constriction allows waters to populate it and that a hydrophilic constriction must be considerably narrower (< 4 Å) than a hydrophobic one to prevent ion permeation (Aryal et al., 2015). These studies align with the observations that mutations of the hydrophobic region of Orai1 create leaky channels (e.g. V102A, F99C) and with molecular simulations that investigated the consequences of these mutations on pore hydration and conduction (Dong et al., 2013; McNally et al., 2012; Megumi Yamashita et al., 2017). Our structures indicate that opening is concerted – the pore of the channel widens along its entire length and includes marked widening of both the hydrophobic and the basic regions. On account of the outward tilting of the subunits, the widening is largest on the cytosolic side of the channel but it also extends to the extracellular side at the selectivity filter.

A theme has emerged from the studies of channels with different architectures that hydrophobic regions within their pores often function as gates that prevent ion permeation when these regions are narrow and permit it when they are widened (reviewed in (Aryal et al., 2015)). Different types of molecular rearrangements engender the dilation and constriction in these “hydrophobic gating” mechanisms. In the superfamily of cation channels that share a pore architecture first observed for a K^+^ channel, which includes voltage-dependent K^+^, Na^+^, and Ca^2+^ channels, a scissor-like movement of helices that also involves helical bending underlies the gating transition of a hydrophobic region of the pore (Jiang et al., 2002b; MacKinnon, 2003). A hydrophobic region in the chloride channel bestrophin opens and closes through a dramatically different mechanism that involves extensive side chain rearrangements throughout the channel (Miller, Vaisey, & Long, 2019). The transition in Orai is mechanistically simpler, involving the outward displacement of subunits through rigid body-like motions as observed in the Orai_H206A_ structure. These examples of mechanisms for dimensional control of the hydrophobic gating regions align with the types of stimuli that control them. Voltage sensor domains in voltage-dependent potassium, sodium and calcium channels are thought to squeeze shut the gates in these channels through mechanical coupling to the scissor-like conformational changes (Jensen et al., 2012; Long, Campbell, & MacKinnon, 2005; Wisedchaisri et al., 2019; Xu et al., 2019). The molecular rearrangements of the gate in bestrophin channels are controlled allosterically by Ca^2+^ and peptide binding to distant sites (G. Vaisey & Long, 2018; George Vaisey, Miller, & Long, 2016). In Orai, the gate opens by outward movements of the subunits and is controlled by STIM from the cytosolic side, where these movements are largest. The structural mechanism for how STIM induces the change is not yet resolved.

## Methods

### Protein expression, purification and cryo-EM sample preparation

The construct used to express Orai_H206A_ is identical to the one previously used (X. Hou et al., 2018). This construct, spanning amino acids 133-341 of *Drosophila melanogaster* Orai, contains a C-terminal YL½-antibody affinity tag (EGEEF), mutations of two non-conserved cysteine residues that improve protein stability (C224S and C283T), and the H206A mutation. The expression of Orai_H206A_ in *Pichia pastoris* and its purification were also as described (X. Hou et al., 2018) with minor modifications. Briefly, lysed cells were re-suspended in buffer (8.3 ml of buffer for each 1 g of cells) containing 150 mM NaCl, 20 mM sodium phosphate, pH 7.5, 0.1 mg/ml deoxyribonuclease I (Sigma-Aldrich), 1:1000 dilution of Protease Inhibitor Cocktail Set III, EDTA free (CalBiochem), 1 mM benzamidine (Sigma-Aldrich), 0.5 mM 4-(2-aminoethyl) benzenesulfonyl fluoride hydrochloride (Gold Biotechnology) and 0.1 mg/ml soybean trypsin inhibitor (Sigma-Aldrich). Lysate was adjusted to pH 8.5 using 1 N KOH while stirring. Next, 0.1 g of n-dodecyl-β-D-maltopyranoside (DDM, Anatrace, solgrade) was added per gram of cells. The sample was stirred at 4 °C for 1 h to extract Orai_H206A_ from the cell membranes. Following extraction, the pH of the sample was adjusted to 7.5 using 1 N KOH. The sample was centrifuged at 30,000 *g* for 45 minutes, and the sample supernatant was filtered using low-protein binding bottle-top filters (Millipore Express Plus 0.22 µm). This filtered supernatant was supplemented with final concentrations of 2 mM EDTA (stock EDTA 200 mM, pH 8.0), 2 mM EGTA (stock EGTA 200 mM, pH 7.0), and 0.2 mM deferoxamine mesylate (Sigma-Aldrich, 20 mM stock in water). YL½ antibody (IgG, expressed from hybridoma cells and purified by ion exchange chromatography) was coupled to CNBr-activated sepharose beads (GE Healthcare) according to the manufacturer’s protocol. Approximately 0.2 ml of beads were added to the sample for each 1 g of *P. pastoris* cells and the mixture was rotated at 4 °C for 1 h. The beads were collected on a column, and washed with 7 column volumes of buffer consisting of 150 mM NaCl, 20 mM sodium phosphate, pH 7.5, 1 mM EDTA, 1 mM EGTA, 0.1 mM deferoxamine mesylate, and 3 mM DDM. Protein was eluted from the beads using a buffer consisting of 150 mM NaCl, 100 mM Tris, pH 8.5, 1 mM EDTA, 1 mM EGTA, 0.1 mM deferoxamine mesylate, 3 mM DDM, and 1 mM EEF peptide (Peptide 2.0). The elutant was concentrated to 500 µl (using an Amicon Ultra 15 100kDa concentrator at 4 °C), filtered using a 0.22 µm filter, and further purified using size exclusion chromatography (Superose 6 Increase 10/300 GL, GE Healthcare) at 4 °C in SEC buffer (150 mM NaCl, 20 mM Tris, pH 8.5, 1 mM DDM). Purified Orai_H206A_ protein fractions were pooled and used to reconstitute into amphipols.

For reconstitution into amphipols, amphipol A8-35 (Anatrace, added as powder) was combined with the purified Orai_H206A_ protein (12:1 wt/wt ratio, amphipol:protein) and the mixture was incubated at 4°C for 14 h. Subsequently, approximately 0.333 g of wet Bio-Beads SM2 (Bio-Rad) were added to the sample for each ml of protein/amphipol mixture. After incubation at 4°C for 5 hours (with rocking), the Bio-Beads were removed by spin filtration and the protein sample was concentrated to 500 µl using an Amicon Ultra 4 (100kDa) concentrator at 4 °C. This amphipol-reconstituted sample was then purified by size exclusion chromatography (Superose 6 Increase 10/300 GL, GE Healthcare) at 4 °C in buffer consisting of 150 mM NaCl, 20 mM Tris, pH 8.5, 0.1 mM deferoxamine mesylate prior to combining it with purified 19B5 Fab.

### Fab production and complex preparation

A monoclonal antibody (designated 19B5) of isotype IgG1 was raised in mice by the Monoclonal Antibody Core Facility of the Memorial Sloan Kettering Cancer Center. The antigen used for immunization was that previously used for X-ray studies: purified Orai (amino acids 133-341 followed by the EEF affinity tag, PDB 6BBH) containing the K163W mutation, which improved protein stability, and has the same overall structure as channel without this mutation (X. Hou et al., 2018; Xiaowei Hou et al., 2012). The antibody selection process included ELISA, western blot, and FSEC analysis (Kawate & Gouaux, 2006) to identify antibodies that bound to native Orai and not SDS-denatured protein. For mapping its binding epitope, experiments included using a construct of Orai that contained an insertion of a Gly-Ala-Gly-Ala sequence in the M1-M2 loop combined with FSEC analysis, which indicated that it bound at or near this loop. We confirmed that 19B5 binds to the extracellular side of the channel by an ELISA-based pull down experiment using intact HEK293 cells that had been transfected with Orai or vector alone. The sequence of the antibody was determined by cDNA sequencing of hybridoma cells (SYD Labs). Intact IgG was expressed using mouse hybridoma cells, purified by ion exchange chromatography and cleaved using papain (1:40 weight-to-weight ratio of papain (Worthington) to IgG) for 3 h at 37 °C to generate the Fab fragment. The Fab fragment was purified using ion exchange chromatography (Mono S, GE Healthcare; using a gradient of 10-500 mM NaCl in 20 mM sodium acetate, pH 5.0), dialyzed into 150 mM NaCl, 20 mM Tris-HCl, pH 8.5, and further purified using size exclusion chromatography (Superose 6 Increase 10/300 GL, GE Healthcare) in SEC buffer (150 mM NaCl, 20 mM Tris, pH 8.5, 0.1 mM deferoxamine mesylate).

Prior to complex formation CaCl_2_ was added to both purified Orai_H206A_ and 19B5 Fab to 5 mM final concentration. The proteins were then combined at a molar ratio of 0.78 : 1 (Fab to Orai monomer, which corresponds to approximately 4.7 Fab molecules per channel assembly), and incubated on ice for 30 minutes. The sample was then concentrated to ∼ 7mg/ml (using a 10K Vivaspin 2 concentrator) and spin-filtered (0.22 µm, Costar). 3 µl of the sample was applied to glow-discharged (10 s) Quantfoil R1.2/1.3 holey carbon grids (Au 400, Electron Microscopy Sciences) and plunge-frozen in liquid ethane using a Vitrobot Mark IV (FEI) robot (settings: 6 °C, blotting time of 2 s, 0 blot force, and 100% instrument humidity). Grids were stored in liquid nitrogen prior to data collection.

### Cryo-EM data acquisition

Clipped grids were loaded into a 300 keV Titan Krios microscope (FEI) equipped with a Gatan K2 Summit direct electron detector (Gatan). Grids were screened first for quality control based on particle distribution, particle density, and the estimated CTF resolution limit across grid regions. Images from the best regions of a single grid were collected at a magnification of 22,500x with a super-resolution pixel size of 0.5442 Å and a defocus range of −0.9 to −2.5 µm. The dose rate was 9 electrons per physical pixel per second, and images were recorded for 10 s with 0.25 s subframes (40 total frames) corresponding to a total dose of approximately 76 electrons per Å^2^.

### Cryo-EM data processing and structure determination

Figure 2 - figure supplement 1 represents the cryo-EM data processing workflow. Movie stacks were dark and gain reference corrected, and subjected to two-fold Fourier cropping to a pixel size of 1.0884 Å. These images were motion corrected and dose weighted using MotionCor2 (S. Q. Zheng et al., 2017). Contrast Transfer Functions for motion-corrected micrographs were estimated using CTFFIND4 (Rohou & Grigorieff, 2015). Micrographs were inspected manually; those with poor-quality features, such as obvious cracks or ice contamination, as well as those micrographs with estimated CTF fits worse than 5 Å were excluded. Out of a total of 4212 collected movies, 3902 passed these two curation criteria. From these micrographs a population of 2,151,623 particles were picked using the Lapacian-of-Gaussian autopicking feature in RELION 3 (Zivanov et al., 2018) and extracted using a particle box size of 384 pixels. 2D classification was conducted in Cryosparc2 (Punjani, Rubinstein, Fleet, & Brubaker, 2017) and classes clearly representing contaminants were excluded, resulting in retention of 1,851,514 particles (86% of the data). These particles were used as input for Cryosparc2 Ab initio 3D model generation, requesting four output models (Figure 2 - figure supplement 1). Particles that yielded Ab initio 3D models that appeared to contain an Orai channel bound to one or more Fabs by visual inspection were subjected to additional rounds of ab initio 3D model generation, from which four distinct models that contained one to three Fab molecules per channel emerged (Figure 2 –figure supplements 1 and 2). Particles containing contain three Fab molecules per channel yielded maps with the highest resolution, exhibited three-fold rotational symmetry (C3), and were selected for all further cryo-EM processing steps. Following Cryosparc2 non-uniform refinement, the selected particles were further sorted by five iterations of heterogeneous 3D classification in Cryosparc2 (using C1 symmetry) to remove remaining assembles that contained fewer than three Fabs or poor particle images; this procedure yielded the final particle set of 85,614 particles. At this point, non-uniform refinement yielded a 3.7 Å reconstruction. The particles were then used for Bayesian polishing in RELION 3. Non-uniform refinement of the polished particles (in cryosparc2) was used to generate the final reconstruction at 3.3 Å. All resolution estimates are based on gold-standard Fourier shell correlation (FSC) calculations.

### Model building and refinement

The atomic model of Orai_H206A_ was manually built into the sharpened cryo-EM map and refined in real space using the COOT software (Emsley, Lohkamp, Scott, & Cowtan, 2010). Previous X-ray structures of Orai (PDB: 4HKR and 6BBH) were used as reference. Further refinement of the atomic model was carried out in PHENIX (Adams et al., 2010) using real-space refinement. The final model has good stereochemistry and good Fourier shell correlation with the map (Figure 2 - figure supplements 3e and 5). Structural figures were prepared with Pymol (pymol.org), Chimera (Pettersen et al., 2004), and HOLE (Smart, Neduvelil, Wang, Wallace, & Sansom, 1996). Electrostatic calculations used the APBS (Baker, Sept, Joseph, Holst, & McCammon, 2001) plugin in Pymol.

### Live cell Ca^2+^ influx measurements

Orai constructs were expressed with N-terminal mCherry tags using a vector that was modified from the pNGFP-EU vector, as described previously (X. Hou et al., 2018; Kawate & Gouaux, 2006). Constructs for wild type Orai (amino acids 120–351) for and full-length *Drosophila melanogaster* STIM (denoted as ‘Orai’ and ‘STIM’ in Figure 1 and Figure 1 - figure supplement 1c-d) have been reported previously (X. Hou et al., 2018). H206A mutation was introduced into the wild type Orai construct (amino acids 120-351) by mutagenesis PCR and is referred to as ‘H206A Orai’.

HEK293 cells were maintained in Dulbecco’s Modified Eagle’s medium (DMEM, the Media Preparation Core of MSKCC) supplemented with 10% fetal bovine serum (Gibco, catalog A31606-02). Approximately 1.5 × 10^6^ cells were co-transfected (in a six-well dish) using the Lipofectamine 3000 transfection reagent (Invitrogen, catalog L3000-015) with 2 µg GCaMP6s plasmid (Chen et al., 2013) and plasmids of Orai and/or STIM constructs, for which the P3000 reagent was used along with Lipofectamine 3000 according to the manufacturer’s protocol (Invitrogen). 0.15 µg Orai-mCherry plasmid and/or 0.6 µg STIM plasmid were used for transfection for samples shown in Figure 1a and Figure 1 - figure supplement 1c. 0.05 µg of plasmid was transfected for H206A Orai and the controls shown in Figure 1b and Figure 1 - figure supplement 1d because larger amounts of H206A Orai plasmid were toxic (presumably due to constitutive activation). Approximately 20 hr after transfection, the cells were trypsinized (Corning, catalog 25–053 Cl), resuspended in FluoroBrite DMEM (GIBCO, catalog A18967-01) supplemented with 2 mM L-glutamine (GIBCO, catalog 25030–081), and seeded to a 384-well plate (∼3×10^4^ cells per well). Approximately 18 hr after seeding, the cells were gently rinsed once with 0Ca solution (10 mM HEPES pH 7.4, 150 mM NaCl, 4.5 mM KCl, 3 mM MgCl_2_, 0.5 mM EGTA and 10 mM D-glucose). After removing the rinse solution, 20 µl of 0Ca solution was added to each well and the cells were incubated for 30–40 min before fluorescence measurements. Fluorescence measurements were recorded using a Hamamatsu FDSS at Ex/Em = 480/527 nm every 2 s throughout the course of the experiment. After recording for 5 min, 10 µl of a 0Ca solution supplemented with thapsigargin (to yield a 1 µM final concentration) and a range of concentrations of purified 19B5 Fab antibody (to yield 0 to 1 µM final concentration) was added. After a subsequent incubation (∼10 min), 10 µl of Ca^2+^-spike solution (10 mM HEPES pH 7.4, 146 mM NaCl, 4.5 mM KCl, 10 mM CaCl_2_ and 10 mM D-glucose) was added to each well to yield a final Ca^2+^ concentration of approximately 2 mM, and recordings were taken for another 8-12 min. The fluorescence traces were generated from an average ±SEM of fluorescence reading from three wells. Fluorescence intensity measurements (F) were plotted directly (Figure 1 - figure supplement 1c,d) or as ΔF (Figure 1) on the Y-axis, where F is the measured fluorescence, F0 is the initial fluorescence value, and ΔF = F-F0.

### Reconstitution and flux assay

Orai constructs were purified and reconstituted into lipid vesicles using published procedures (X. Hou et al., 2018; Xiaowei Hou et al., 2012). The constructs (amino acids 133-341 of Orai with P288L, H206A or V174A mutations) are identical to one another except for the indicated mutations (X. Hou et al., 2018; Xiaowei Hou et al., 2012). Briefly, a lipid mixture containing 15 mg/ml POPE (1-palmitoyl-2-oleoyl-sn-glycero-3-phosphocholine) and 5 mg/ml POPG (1-palmitoyl-2-oleoyl-sn-glycero-3-phospho(1’-rac-glycerol)) was prepared in water and solubilized with 8% (w/vol) n-octyl-β-D-maltopyranoside (V174A and P288L) or 8% (w/vol) n-decyl-β-D-maltopyranoside (H206A). Purified Orai protein was mixed with the solubilized lipids to obtain a final protein concentration of 0.1 mg/ml and a lipid concentration of 10 mg/ml. Detergent was removed by dialysis (15 kDa molecular weight cutoff) at 4 °C for 5-7 days against a reconstitution buffer containing 10 mM HEPES pH 7.0, 150 mM KCl and 0.2 mM ethylene glycol tetraacetic acid (EGTA), with daily buffer exchanges and utilizing a total volume of 14 l of reconstitution buffer. The reconstituted sample was sonicated (∼30 s), aliquoted, flash-frozen in liquid nitrogen and stored at −80 °C.

The fluorescence-based flux assay was performed as previously described (X. Hou et al., 2018; Xiaowei Hou et al., 2012). In brief, vesicles were rapidly thawed (using 37 °C water bath), sonicated for 5 sec, incubated at room temperature for 2-4 hours before use, and then diluted 100-fold into a flux assay buffer containing 150 mM n-methyl-d-glucamine (NMDG), 10 mM HEPES pH 7.0, 0.2 mM EGTA, 0.5 mg/mL bovine serum albumin and 0.2 µM 9-amino-6-chloro-2-methoxyacridine (ACMA, from a 2 mM stock in DMSO). Fluorescence intensity measurements were collected every 30 sec on a SpectraMax M5 fluorometer using Softmax Pro 5 software (Molecular Devices; excitation and emission set to 410 nm and 490 nm, respectively). The proton ionophore carbonyl cyanide m-chlorophenyl hydrazine (CCCP, 1 µM from a 1 mM stock in DMSO) was added between the 150 and 180 sec time points and the sample was mixed by pipette. The potassium ionophore valinomycin (2 nM from a 2 µM stock in DMSO) was added near the end of the experiment to establish baseline fluorescence and confirm vesicle integrity.

## Data availability

The electron microscopy map and the atomic model have been deposited in the Electron Microscopy Data Bank (EMDB) and in the Protein Data Bank (PDB) with deposition ID D_1000251386.

## Acknowledgments

We thank Richard Hite and members of the laboratory for discussions, M.J. de la Cruz of the Memorial Sloan Kettering Cancer Center Cryo-EM facility for help with data collection, and Frances Weis-Garcia and the staff of the Antibody and Bioresource core facility at Memorial Sloan Kettering Cancer Center. This work was supported by NIH grants R01GM094273 and R35GM131921 (to S.B.L.) and a core facilities support grant to Memorial Sloan Kettering Cancer Center (P30CA008748).

## Author Contributions

X.H and I.R.O. performed cryo-EM and associated studies. X.H. performed functional analyses. L.P. assisted with antibody development. S.B.L. directed and contributed to the research. All authors contributed to data analysis and the preparation of the manuscript.

## Author Information

The authors declare no competing financial interests. Correspondence and requests for materials should be addressed to S.B.L.

---

**Figure 1 – figure supplement 1.**
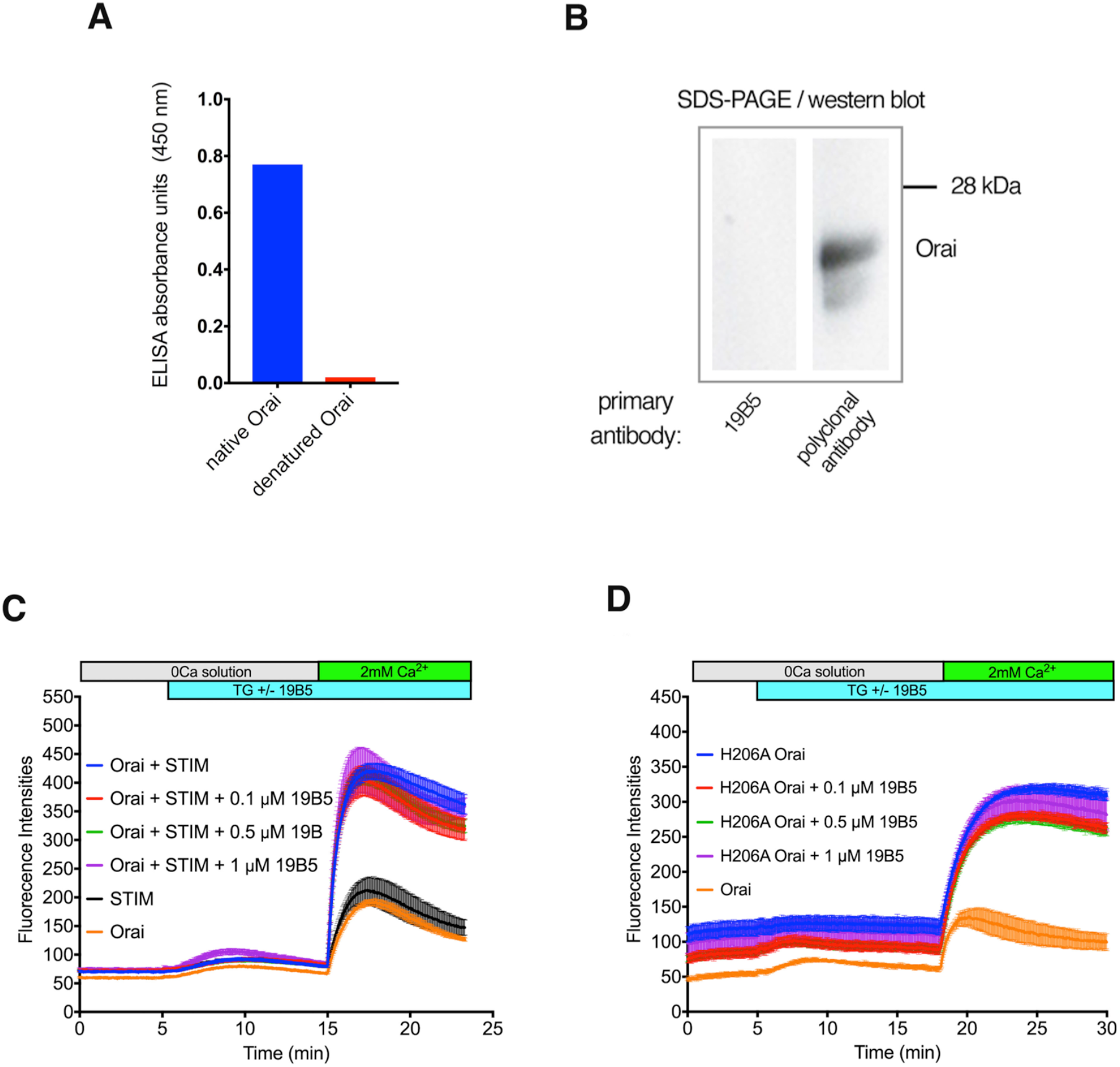
Further analyses of the 19B5 antibody. **a**, ELISA analysis indicates that 19B5 preferentially binds to native rather than denatured Orai. An ELISA assay was performed using plates that had been coated with purified Orai (native Orai) or with purified Orai that had been denatured using SDS (denatured Orai). Absorbance (at 450 nM) indicates binding of 19B5. **b**, Western blot analysis of 19B5, showing that 19B5 does not recognize denatured Orai. A polyclonal antibody against Orai (obtained from the mouse used to generate 19B5) contains antibodies that recognize denatured Orai protein (∼ 24 kDa), and is shown as a control. **c and d**, Raw data for the Ca^2+^ influx measurements shown in Figure 1, with fluorescence intensity on the *y*-axis. The variable initial intensity value for H206A Orai (d) can be attributed to the constitutive activity of this mutant.

**Figure 2 – figure supplement 1.**
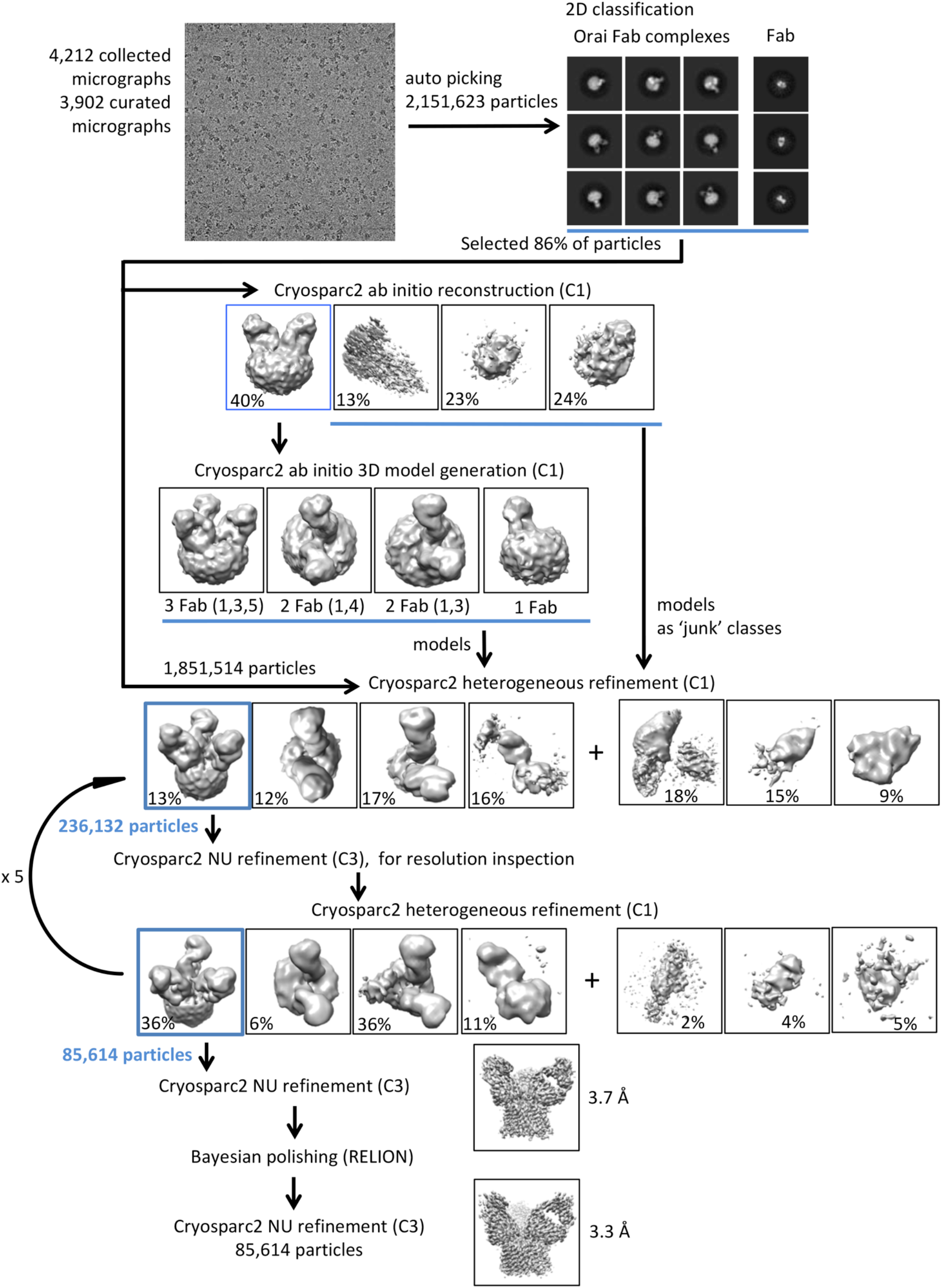
Flowchart for cryo-EM data processing of the Orai_H206A_-Fab complex. Details can be found in the Methods.

**Figure 2 – figure supplement 2.**
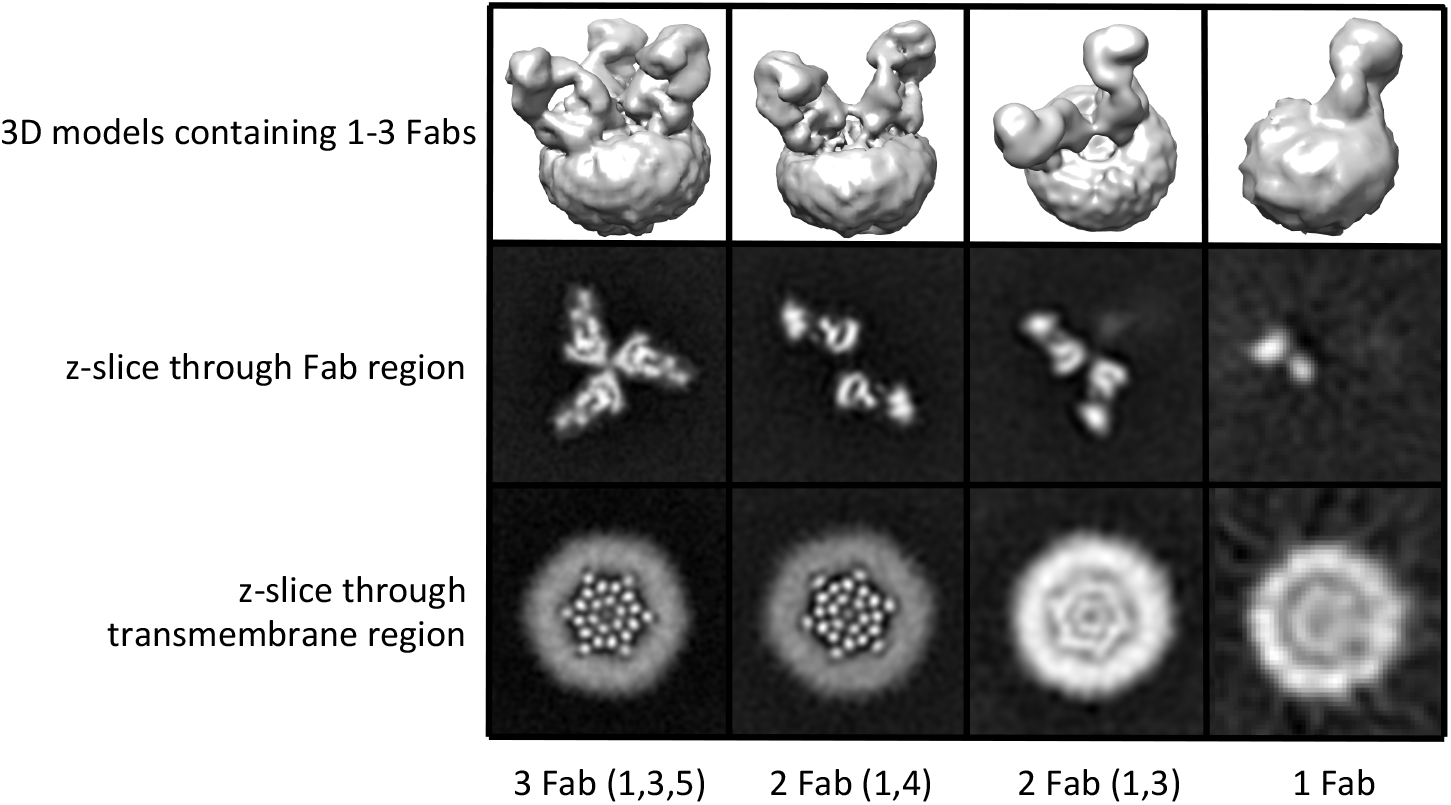
3D reconstructions of Orai_H206A_-Fab complexes containing between one and three Fab molecules. 3D reconstructions of complexes containing between one and three Fabs are shown in the upper panels (Chimera representations). Slices of the Fab and transmembrane portions of the 3D reconstructions are shown in the middle and lower panels, respectively (the views are from the extracellular side these panels). The halo in the lower panels is due to the amphipols surrounding the transmembrane portion of Orai. Complexes containing two Fab molecules have Fabs arranged in two alternative configurations (1,4) or (1,3).

**Figure 2 – figure supplement 3.**
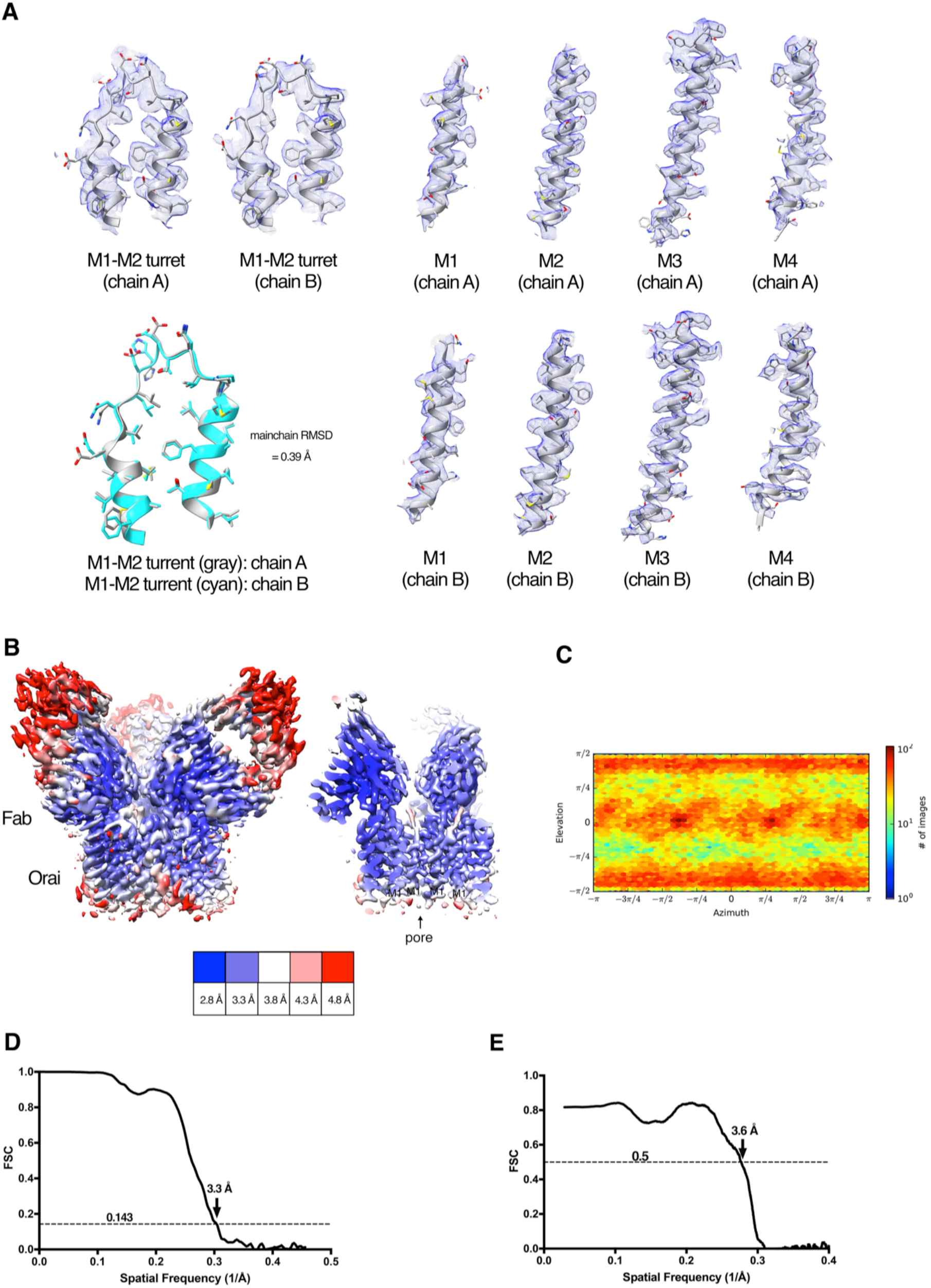
Cryo-EM structure determination and density. **a**, Densities (blue mesh) of indicated regions of Orai in the Orai-Fab complex are shown in the context of the atomic model (cartoon and stick representation). A superposition of the M1-M2 turrets from two neighboring subunits is also shown (lower left panel, main chain RMSD=0.39 Å). **b**, Local resolution of the map estimated using the Local Resolution Estimation function of Cryosparc2 (colored as indicated). An overall view (left panel) and a cross-section (right panel) are shown. **c**, Angular orientation distribution of the particles used in the final reconstruction. **d**, Gold-standard FSC curve of the final 3D reconstruction. The resolution is 3.3 Å at the FSC cutoff of 0.143. **e**, FSC curve showing the correlation of the atomic model and cryo-EM map.

**Figure 2 – figure supplement 4.**
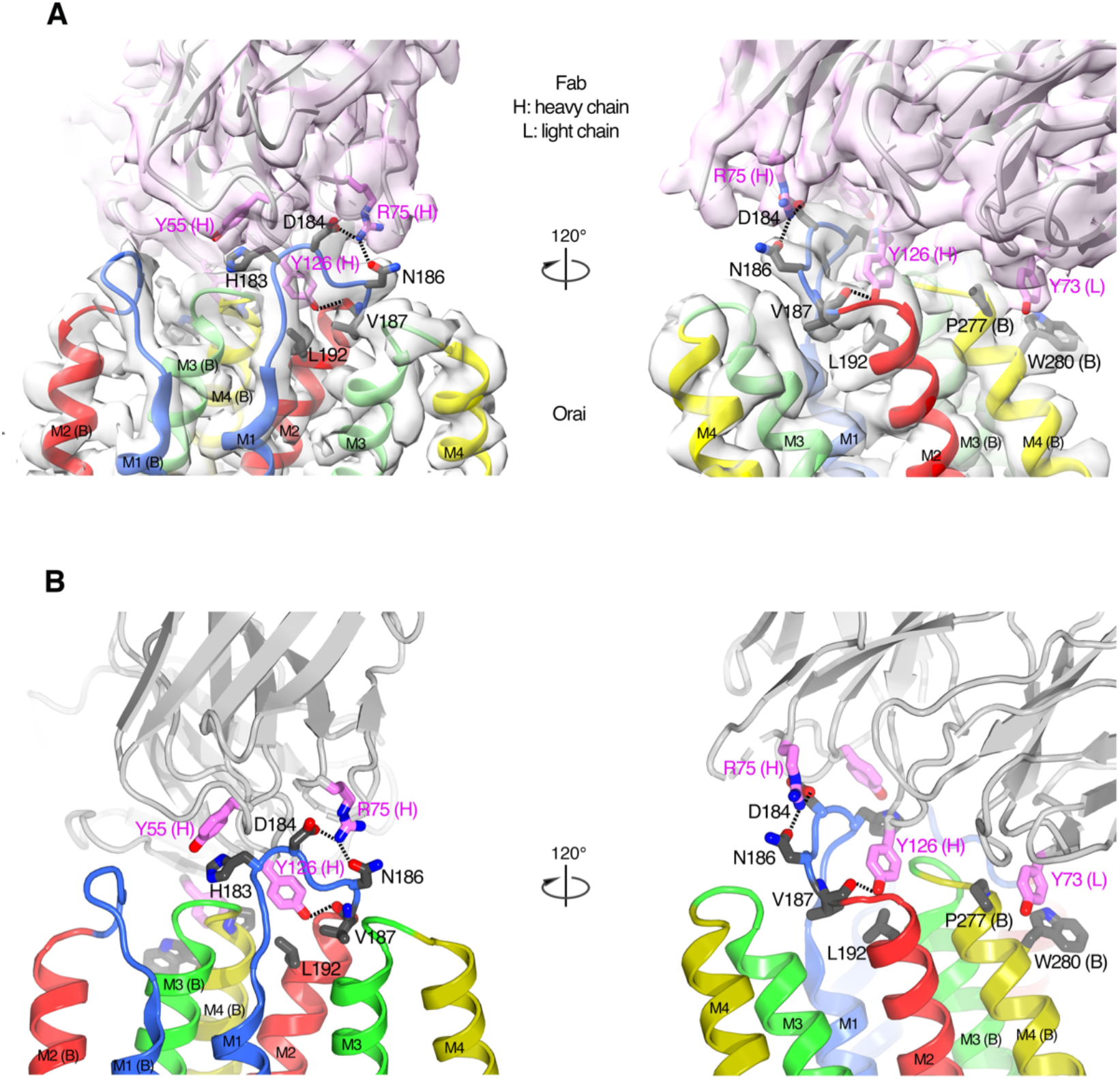
The Orai-Fab interface. **a**, Cartoon representations of the antibody (violet) and two neighboring subunits of Orai (labeled A and B and colored according to transmembrane α-helices as in Figure 2) are shown with the corresponding cryo-EM density (semitransparent surfaces). Amino acids in the Fab-Orai binding interface are drawn as sticks (with carbon atoms colored violet for the Fab and gray for Orai, and with oxygen and nitrogen atoms colored red and blue, respectively). Dashed lines indicate hydrogen bonds. Two perspectives are shown (left and right panels). **b**, Analogous depictions as in (a), with the density removed for clarity.

**Figure 2 – figure supplement 5.**
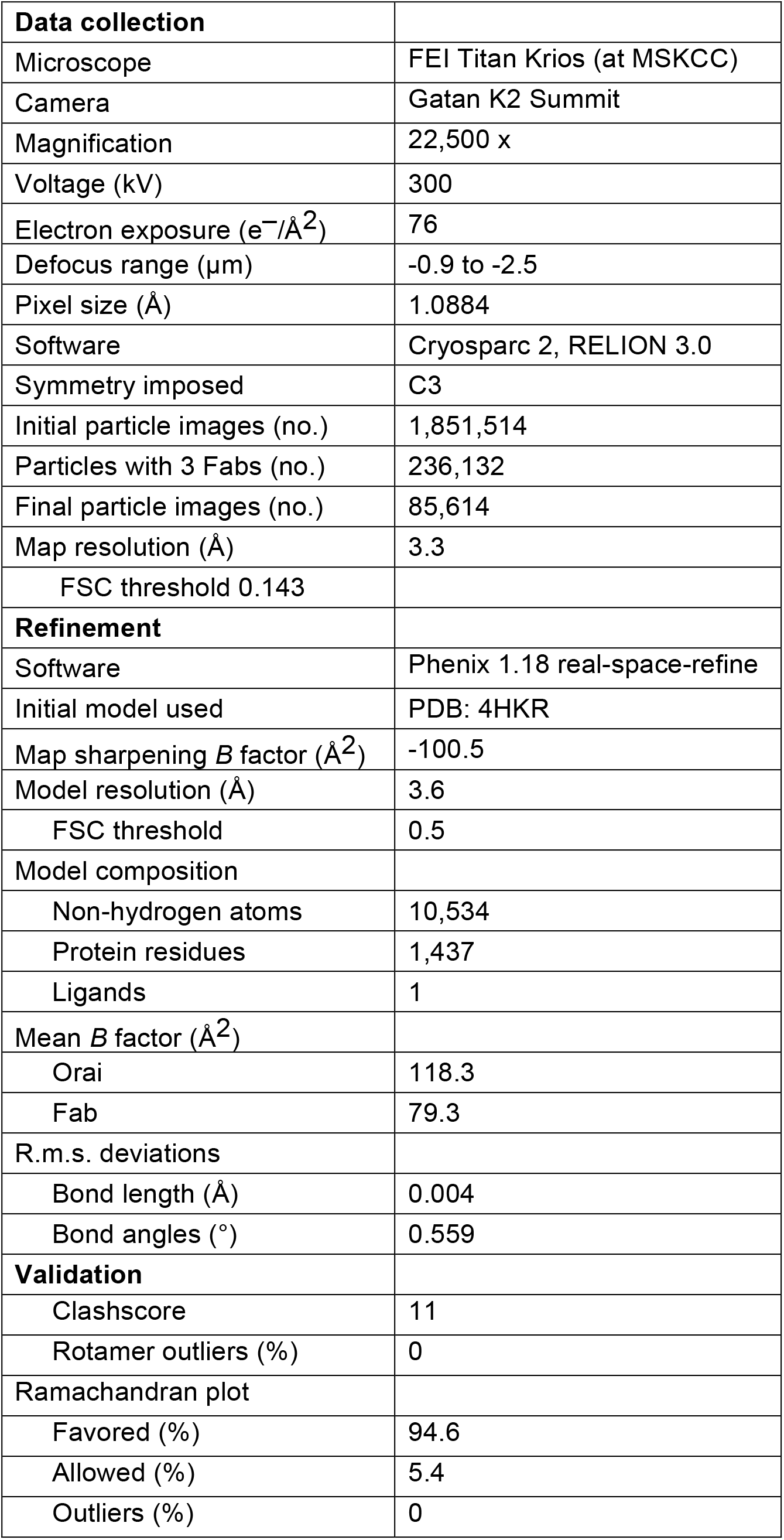
Cryo-EM data collection, refinement and validation statistics.

**Figure 9 – figure supplement 1.**
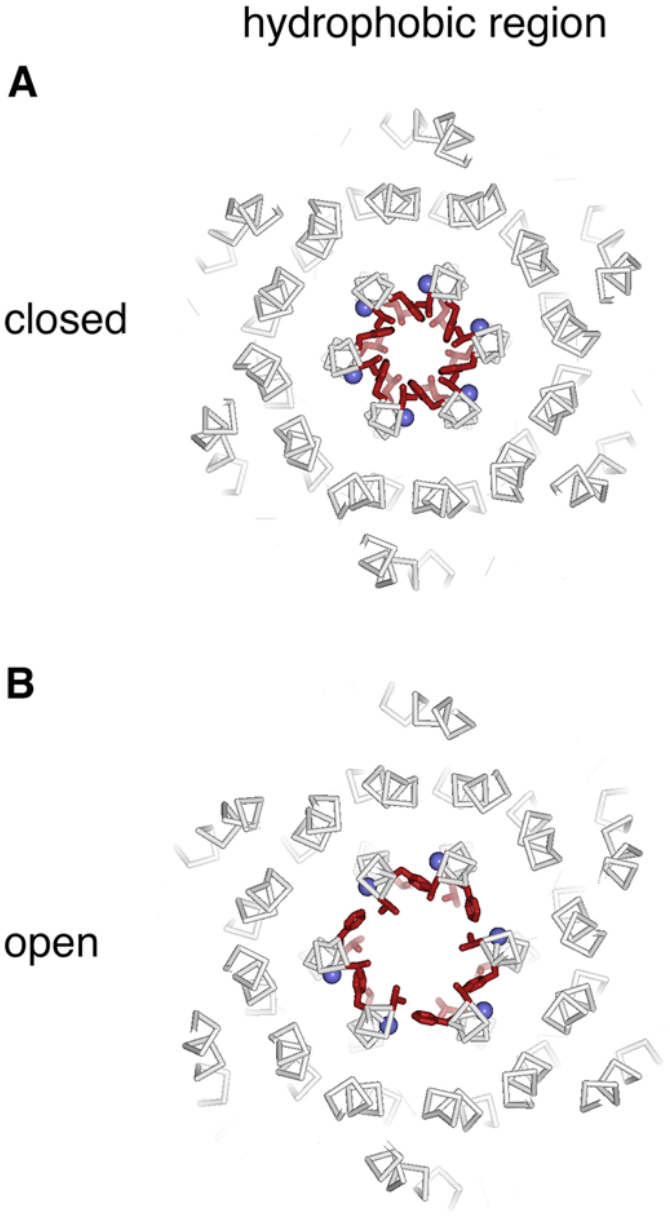
Hydrophobic region. **a**,**b**, Slices analogous to Figure 9, showing a slab containing the entire hydrophobic region of each pore.

**Figure 9 – figure supplement 2.**
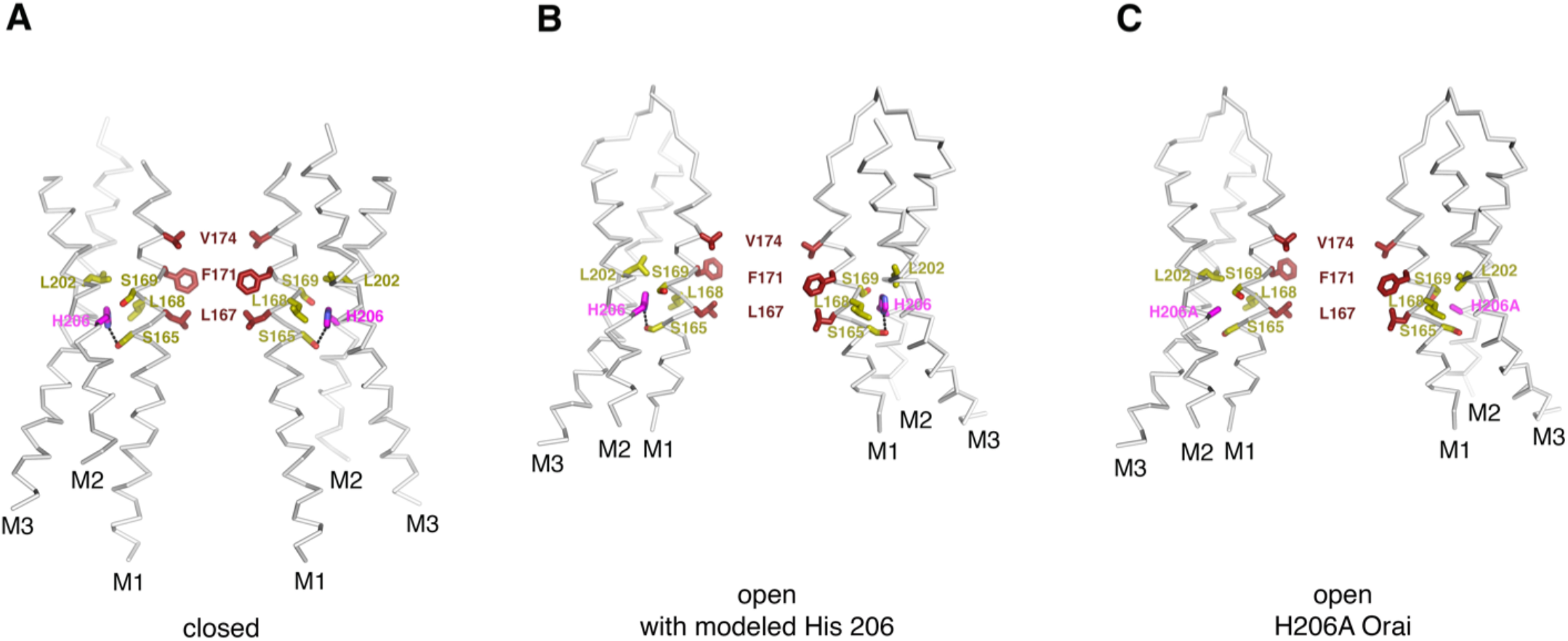
Residue 206 in the closed and open conformations. **a**, In the closed structure (PDB ID:4HKR), His 206 forms a hydrogen bond with Ser 165 (dashed line) and is in van der Waals contact with Leu 202, Leu 168, and Ser 169. **b**, Modeling of a histidine residue at position 206 into the Orai_H206A_ open structure. The side chain of His 206 can be accommodated in the cryo-EM structure of Orai_H206A_ with minor adjustments to the surrounding amino acid side chains (refer to panel c for these adjustments). In the model, and in a similar manner as in the closed structure, His 206 forms a hydrogen bond with Ser 165 (dashed line) and is in van der Waals contact with Leu 202, Leu 168, and Ser 169. **c**, Depiction of the structure of Orai_H206A_. In this open structure, the alanine at residue 206 eliminates the hydrogen bond to Ser 165 but the H206A residue still interacts with Leu 168 and Leu 202. The structures in (a-c) are drawn as ribbons with the indicated residues shown as sticks. Residue 206 is colored magenta; residues that are in close contact with it are colored yellow; and the residues of M1 that form the hydrophobic region of the pore are colored dark red.

**Figure 9 – figure supplement 3.**
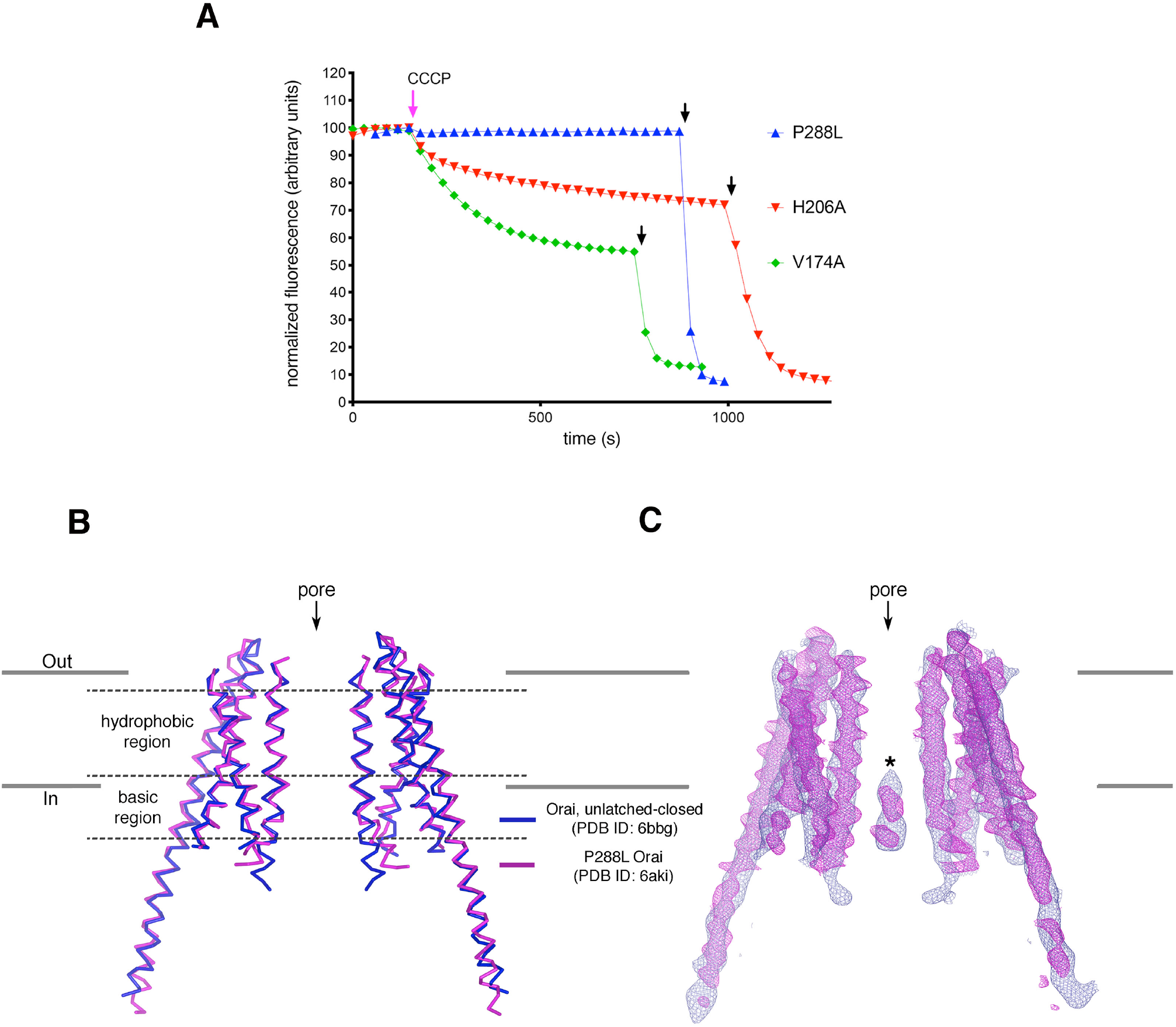
Analyses of P288L Orai. **a**, Ion flux measurements of purified Orai mutants. K^+^ flux measurements for Orai with the H206A, P288L, or V174A mutations under divalent-free conditions. The assay was performed as previously described (X. Hou et al., 2018; Xiaowei Hou et al., 2012). Proteoliposomes containing purified Orai with the indicated mutations were loaded with 150 mM KCl and were diluted 50-fold into flux buffer containing a fluorescent pH indicator (ACMA) and 150 mM N-methyl-D-glucamine (NMDG) to establish a K^+^ gradient (Methods). After stabilization of the fluorescence signal (150 s), a proton ionophore (CCCP) was added. An electric potential arising from K^+^ efflux drives the uptake of protons, which quenches the fluorescence of ACMA. The time-dependent decrease in fluorescence observed for the H206A and V174A after the addition of CCCP is indicative of K^+^ flux, whereas K^+^ flux is not detected for the P288L mutant. Valinomycin, which renders the vesicles permeable to K^+^, was added near the end of the experiment (black arrows) as a positive control and to establish baseline fluorescence. Traces were normalized by dividing by the initial fluorescence value, which was within ±10% for each experiment. The experiments shown used a protein-to-lipid ratio of 1:100 (wt/wt). Higher concentrations of the P288L mutant (ratios of 1:10 and 1:50) yielded analogous results; ion flux was not detected through P288L Orai. As expected, wild type Orai does not show K^+^ flux in this assay as it is not active without STIM (X. Hou et al., 2018; Xiaowei Hou et al., 2012). **b**, Superposition of an X-ray structure of P288L Orai (Liu et al., 2019) with an X-ray structure of Orai without this mutation (WT Orai) in the unlatched-closed conformation, in which the pore is closed (X. Hou et al., 2018). Two opposing subunits from each structure are shown, with the WT and P288L structures depicted as blue and magenta ribbons, respectively. The RMSD of the superposition is 1.5 Å (for Cα atoms from M1 to M4). **c**, Superposition of the electron densities. Electron densities for WT and P288L Orai are drawn in blue and magenta mesh representations, respectively, covering the two subunits shown in (b). The map of WT Orai (contoured at 1.3 σ) was calculated from 20 to 6.9 Å resolution using amplitudes and MR-SAD phases that had been improved by NCS averaging, solvent flattening and histogram matching (as described in (X. Hou et al., 2018)). The density from the P288L Orai structure (2Fo-Fc electron density map, contoured at 1.5 σ) was generated using map coefficients FWT and PHWT downloaded from RCSB Protein Data Bank (PDB ID: 6AKI). The structure and electron density superpositions show that the two structures are indistinguishable within the transmembrane region of the channel and that both structures contain the narrow hydrophobic region characteristic of a closed pore. An asterisk indicates the density assigned to an anion/iron complex that was observed in the basic region of the closed pore (X. Hou et al., 2018; Xiaowei Hou et al., 2012). Similar density is present in the P288L structure, but this density was modeled as Cl^-^ (Liu et al., 2019). However, because Cl^-^ ions would not be visible in low-resolution maps, we suggest that the density more likely represents the anion/iron complex that is observed when the pore is closed (X. Hou et al., 2018; Xiaowei Hou et al., 2012).\

**Video 1**. Opening transition, depicted as in Figure 9, showing a morph between the closed and open conformations.

**Video 2**. Opening transition, showing a side view of the pore. Depicted similarly to Figure 9, this video shows a side view of a morph between the closed and open conformations. Two opposite subunits are drawn as ribbons with amino acids in the hydrophobic region of the pore depicted as red sticks. Gly 170 residues are shown as blue spheres.

